# Patient brain organoids identify a link between the 16p11.2 copy number variant and the *RBFOX1* gene

**DOI:** 10.1101/2021.11.21.469432

**Authors:** Milos Kostic, Joseph J. Raymond, Beata Henry, Tayfun Tumkaya, Jivan Khlghatyan, Jill Dvornik, Jack S. Hsiao, Seon Hye Cheon, Jonathan Chung, Yishan Sun, Ricardo E. Dolmetsch, Kathleen A. Worringer, Robert J. Ihry

**Affiliations:** Neuroscience, Novartis Institutes for BioMedical Research, Cambridge, MA, USA; Chemical Biology and Therapeutics, Novartis Institutes for BioMedical Research, Cambridge, MA, USA

## Abstract

Copy number variants (CNVs) that delete or duplicate 30 genes within the 16p11.2 genomic region give rise to a range of neurodevelopmental phenotypes with high penetrance in humans. Despite the identification of this small region, the mechanisms by which 16p11.2 CNVs lead to disease are unclear. Relevant models, like human cortical organoids (hCOs), are needed to understand the human-specific mechanisms of neurodevelopmental disease. We generated hCOs from 18 patients and controls, profiling 167,958 cells with single cell (sc)RNA-seq. Analysis revealed neuronal-specific differential expression of genes outside of the 16p11.2 region that were related to cell-cell adhesion, neuronal projection growth, and neurodevelopmental disorders. Furthermore, 16p11.2 deletion syndrome organoids exhibited reduced mRNA and protein levels of RBFOX1, a gene which can also harbor CNVs linked to neurodevelopmental phenotypes. We found that many genes previously shown to be regulated by RBFOX1 are also perturbed in organoids from patients with 16p11.2 deletion syndrome, and thus identified a novel link between independent CNVs associated with neuronal development and autism. Overall, this work suggests convergent signaling, which indicates the possibility of a common therapeutic mechanism across multiple rare neuronal diseases.

## INTRODUCTION

Copy number variants (CNVs) play a major role in the etiology of neuropsychiatric and neurodevelopmental disorders. A CNV may reside in a single gene, such as *RBFOX1,* which encodes a protein regulating mRNA alternative splicing in neurons and is located on chromosome 16p13.3 ^1^. *RBFOX1* CNVs are linked to autism spectrum disorder (ASD), intellectual disability (ID) and epilepsy ^1–4^. A CNV may also span a specific chromosomal segment or cytogenetic band, such as 1q21.1, 7q11.23, 15q11.2, 16p11.2, 17q12, and 22q11.2 (chromosome number, p/q denotes long/short arm of each chromosome, band number), causing deletion or duplication of a whole set of resident genes ^5, 6^. The CNVs spanning chromosome 16p11.2 have been associated with multiple neuropsychiatric and neurodevelopmental disorders ^7^ and are the focus of our current study. Some clinical diagnoses or phenotypes are common to both 16p11.2 hemideletion (1 copy loss) and hemiduplication (1 copy gain), such as autism spectrum disorder (ASD), intellectual disability (ID), and epilepsy ^7–14^. Other diagnoses or phenotypes are unique to the actual copy number and can be reciprocal between hemideletion and hemiduplication. For example, 16p11.2 hemideletions are linked to macrocephaly and obesity, while hemiduplications are linked to microcephaly and low body mass index (BMI) ^15, 16^. Furthermore, clinical phenotypes may also vary by the exact chromosomal breaking points of a 16p11.2 CNV. In humans, the most common genomic breakpoints (BPs) underlying either 16p11.2 hemideletion or hemiduplication are BP4-BP5, which straddle a ∼600 kilobase pair (kbp) centromere-proximal region of 16p11.2 ^17, 18^. However, some individuals carry CNVs with BPs flanking a more distal region (e.g. BP1-BP3, BP2-BP3, BP1-BP4) or flanking both proximal and distal regions (e.g. BP1-BP5). Interestingly, susceptibility to schizophrenia was only associated with proximal duplications and distal deletions, suggesting differential roles among resident genes and/or non-coding elements between the proximal and distal regions of 16p11.2 in human brain development and function ^19, 20^. In addition, hemiduplications, but not hemideletions, are associated with schizophrenia (SCZ) ^18, 21^.

Human brain development and behavior are different in rodents, and thus rodent models are insufficient to fully understand human disease ^22, 23^. The species divergence is particularly relevant for 16p11.2 CNVs, because mice genetically modified for a chromosomal segment homologous to human 16p11.2 exhibit dissonant phenotypes relative to humans. For example, the hemideletion mice showed microcephaly and low BMI, whereas the hemiduplication mice had high BMI ^24, 25^. With the advent of the ability to reprogram human somatic cells into induced pluripotent stem cells (iPSCs) ^26, 27^, we can now generate *in vitro* neuronal models for studying human disease mechanisms ^28–30^. Previous studies have used two-dimensional (2D) iPSC neuronal models to examine the effects of 16p11.2 hemideletion and hemiduplication, and identified synapse, neurite, and dendrite length phenotypes, as well as transcriptional disruptions of genes outside of the 16p11.2 region ^31, 32^. Recently, three-dimensional (3D) hiPSC-derived brain organoids have been used to improve the experimental modeling of brain development in physiological and diseased states ^33–37^. With the advent of single cell RNA sequencing (scRNA-seq) ^38^, it is now feasible to profile transcriptional landscapes of brain organoids with cellular resolution ^37, 39–42^.

We aimed to better understand the disease mechanisms and affected cell types in 16p11.2 patients by leveraging human cortical organoid (hCO) models. To this end, we established a semi-automated protocol to generate and deeply characterize the cellular composition and cell-specific gene expression of hCOs by performing scRNA-seq at three differentiation time points. These data were integrated with a fetal brain atlas and we observed that our hCO model recapitulates crucial aspects of neurogenesis. We performed a large-scale guided differentiation of hCOs from 18 human iPSC lines derived from controls and patients with 16p11.2 hemideletion or hemiduplication. Using scRNA-seq as a readout, we observed reproducible cellular composition within and across hCO lines. Importantly, we revealed cell type-specific changes in gene expression between hemideletion patients and controls. These differentially expressed (DE) genes are important for neurite formation, neuronal cell-cell adhesion, and synaptic processes, and have been previously associated with neurodevelopmental disorders. Furthermore, the reduction in RBFOX1, an RNA binding protein within ASD-associated CNVs, was validated at the protein level. Consistent with this, RBFOX1 target mRNAs were disrupted in 16p11.2 patient organoids. Together these data suggest that cell type-specific changes in gene expression may contribute to developing a 16p11.2 pathogenic phenotype and that independent rare CNVs and resident genes may exhibit effects through convergent molecular signaling pathways.

## RESULTS

### Building a platform for robust human cortical organoid generation

We modified existing protocols using 3 patterning factors, with optimized doses and duration, to differentiate cells to neuronal lineages from human iPSCs into hCOs (see Methods) ^36, 43^ (Figure 1A). To scale up and minimize human error, we utilized an automated liquid-handling system (Hamilton STAR) during the most demanding part of the hCO differentiation protocol (until day 25) (Figure 1A) (see Methods). Using single cell transcriptomics, we aimed to compare a large-scale cohort of hCOs at different timepoints and batches. To reduce the technical challenges associated with working with many patient hCOs, we uncoupled the requirement of dissociating hCOs and performing scRNA-seq the same day. Methanol fixation of dissociated hCOs was optimized to enable long term sample storage and the ability to rerun control samples during scRNA-seq library construction to control batch effects and monitor reproducibility ^44^ (Figure 1A).

**Figure 1.**
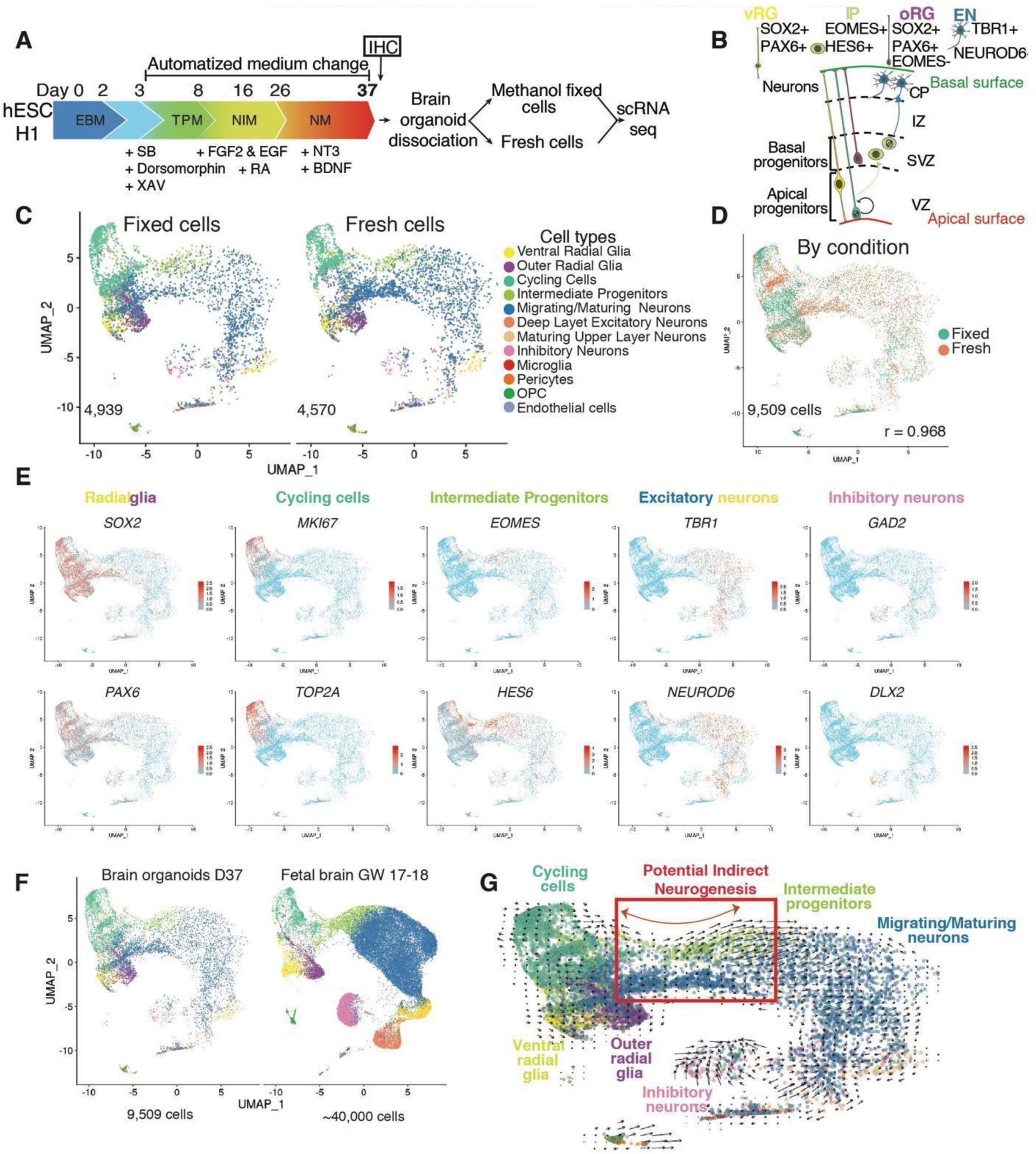
Semi-Automated Generation of Human Cortical Organoids Yields Crucial Aspects of Human Neurogenesis. (A) Schematic of human cortical organoids (hCOs) differentiation and methanol fixation protocols, cell line hESC-H1. (B) Schematic of neocortical development and its main cell types. (C) Uniform Manifold Approximation and Projection (UMAP) embedding plot of 4,939 methanol fixed and 4,570 fresh cells, day 37 hCOs, with all identified cell types. (D) UMAP plot showing overlap of methanol-fixed and fresh cortical organoid cells, Pearson’s correlation r = 0.968. (E) UMAP plots showing expression of canonical gene markers for main cell types which were identified in fresh and fixed cells combined at day 37. (F) UMAP plots integration of hCOs (day 37, 9,509 fixed and fresh cells) with fetal brain atlas (GW17-18, ∼40,000 cells, ^47^), majority of hCOs cells is allocated to the neuronal progenitor and immure neurons identities; GW – gestational week. (G) RNA velocity is projected onto a predefined UMAP plot, hCOs (day 37) fixed and fresh cell combined. Length of the arrow annotates the transcriptional dynamics, and the direction of the arrow points to the future state of cells. Inset, revealing the potential indirect neurogenesis in hCOs. EBM, Embryoid body medium; TPM, Telencephalon patterning medium; NIM, Neural induction medium; NM, neuronal medium; vRG, ventral radial glia; oRG, outer radial glia; IP, intermediate progenitors; EN, excitatory neurons; Ventricular Zone, VZ; Subventricular Zone, SVZ; Intermediate Zone, IZ; Cortical Plate, CP.

First, cellular diversity of hCOs was characterized under both fresh and fixed conditions by splitting a sample of twenty (10 fresh and 10 fixed) dissociated hCOs on day 37 of differentiation. Freshly dissociated and fixed cell suspensions were compared with scRNA-seq ^45^ (Figure 1A). In total, we profiled 9,508 cells: 4,939 fixed and 4,570 fresh cells. We assessed cell quality, where cells with less than 5% of mitochondrial reads are judged to be high quality ^46^. The quality was similar for each condition, exhibiting less than 5% of reads aligning to mitochondrial transcripts (Figure S1A). We used Seurat’s TransferAnchors function (see Methods) to identify the putative cell types present in hCOs (at day 37) by comparing them to fetal brain cells from a reference dataset ^47^. Both fresh and fixed hCOs contained all the main types of neuronal progenitors found in fetal brain samples – ventral and outer radial glia (vRG, oRG; markers *SOX2, PAX6*), Cycling Cells (CycCells; markers *MKI67, TOP2A*), intermediate progenitors (IPs; markers *EOMES, HES6*), and different types of excitatory neurons including migrating/maturing neurons (MigNs, MatN), deep and upper layer excitatory neurons (DlExNs, UlNs), expressing markers *TBR1* and *NEUROD6* (Figures 1B, 1C and 1E). Interestingly, we found a small population of inhibitory neurons (INs; markers *GAD2, DLX2*), similar to previous reports ^37, 41^ (Figures 1B, 1C, and 1E). In addition, the majority of day 37 hCO cells were progenitor cell types, with a smaller population of mature neurons (Figure 1F). Lastly, we found a minor population (< 4% of total cells number) of oligodendrocyte precursor cells (OPCs), microglia, pericytes, and endothelial cells. Overall, we found that methanol fixation does not significantly alter the transcriptional profile and cellular composition (Figures 1C, 1D, and S1B).

To analyze the lineage relationship between the predominant cell types, we performed RNA-velocity analysis using velocyto ^48^. The 10X Genomics scRNA-seq platform captures both unspliced and spliced mRNA. RNA-velocity takes advantage of this by examining the relative abundance ratio of unspliced and spliced mRNA. This is used to estimate transcriptional changes and cell lineage direction. The most prominent lineage was originating from neural progenitor cells, transiently crossing the IPs and differentiating into excitatory neurons (Figures 1G; red box). This suggests hCOs undergo indirect neurogenesis ^49–53^, where neural progenitor cells undergo asymmetric division to give rise to IPs, which eventually differentiate into excitatory cortical neurons. Together, these results show the semi-automated platform for generating hCOs yields cells which model human embryonic neurogenesis.

### Characterizing the maturation of human cortical organoids

To benchmark cellular diversity of hCOs during maturation, we performed scRNA-seq on 30-, 90- and 166-day old hCOs (Figures 2A-2D). We methanol fixed cells at each time point, which allowed us to process all samples for scRNA-seq at the same time to avoid batch effects. In total, we profiled 24,662 cells and identified 13 different cell types, of which 9 of the following cell types were predominant (∼98.55% of total population): vRGs, oRGs, CycCells, IPs, MigNs, MatNs, DlExNs, UlNs, and INs (Figures 2B, 2C, and S1C). At day 30, more than 50% of all cells were progenitors: vRG, oRG, CycCells, and IPs. By day 90, progenitor populations decreased to under 20%, and by day 166, vRG virtually disappeared. These changes were concomitant with a dramatic increase in neuronal populations (Figures 2B, 2C, and S1C). Similar to previous experiments (Figure 1C), we detected INs expressing *DLX5* and *GAD2*, and the fraction of INs peaked at D90 (Figures 2C and 2E). The fraction of oRGs (*PTPRZ1*, *HOPX*) increased from 1% (D30) to 10% at day 90 and then decreased to 3% by day 166, which reflects human neurogenesis where peak of oRG production is in mid-gestational period of cortical development ^54–56^. Additionally, 12% of cells at day 166 were identified as maturing upper layer excitatory neurons expressing (Figure 2C and 2E). This is consistent with temporal layer-specific patterning of neurons, with DlExNs being generated first in early neurogenesis, and UlNs generation peaking in mid to late neurogenesis (Figures 2B-2D) ^57–60^.

**Figure 2.**
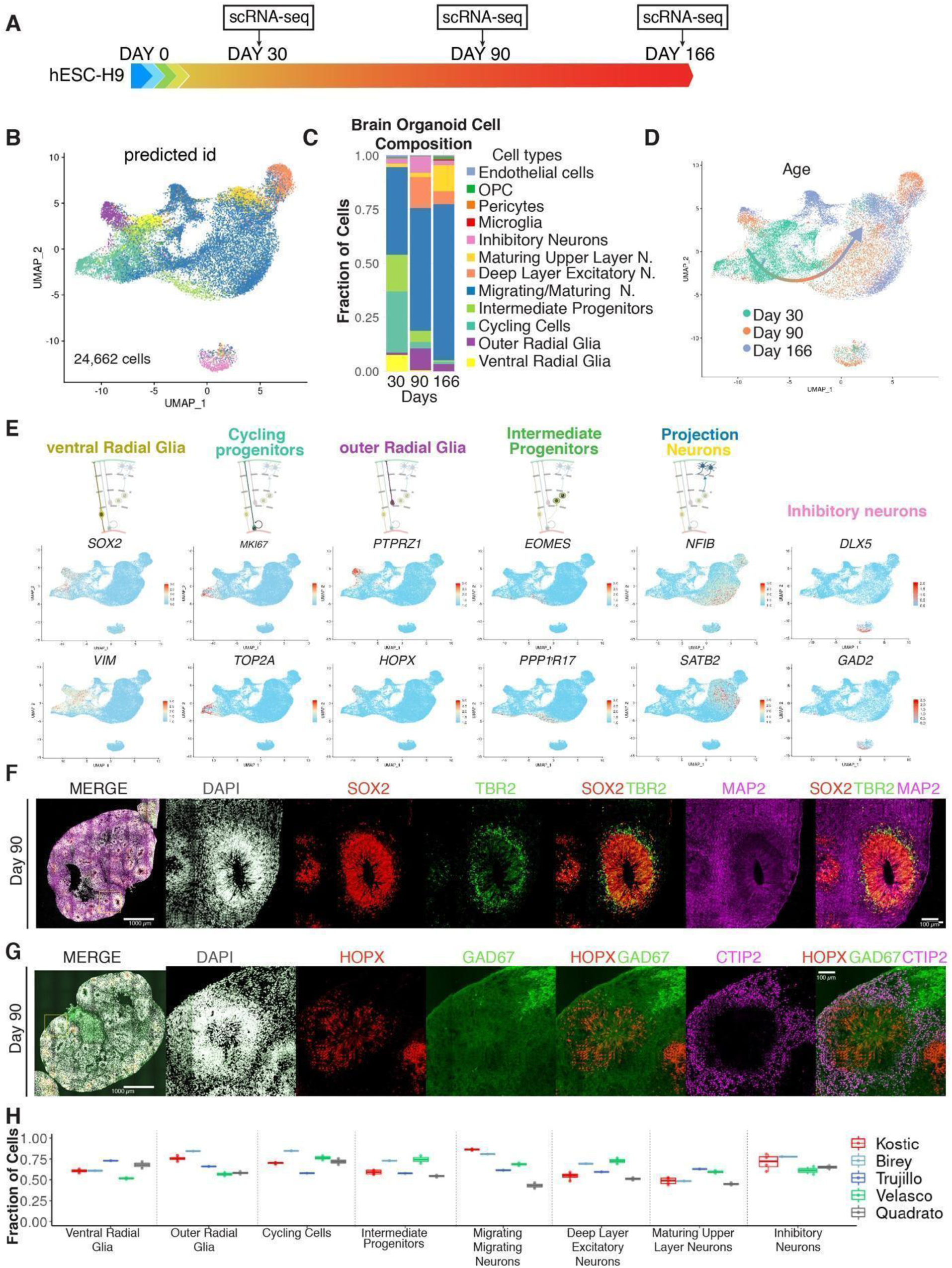
Longitudinal scRNA-seq Profiling of Human Cortical Organoid Maturation. (A) Timeline of hCOs differentiation, cell line hESC-H9. (B) scRNA-seq dataset generated from three different ages of hCOs (Day 30, Day 90, Day 166, total - 24,662 cells). (C) Stacked bar plot showing relative distribution of cell types in hCOs measured by scRNA-seq split by age Day 30, Day 90, and Day 166, each column represents 10-20 randomly sampled organoids pulled and analyzed. (D) UMAP plot of hCOs scRNA-seq dataset split by age (Day 30 [green] - 8,955 cells, Day 90 [orange] - 5,969 cells, and Day 166 [blue] - 10,011 cells). (E) UMAP plots showing expression of markers for main types of cells identified in hCOs, vRG (*SOX2*, *VIM*), cycling progenitors (*MKI67*, *TOP2A*), oRG (*PTPRZ1*, *HOPX*), IP (*EOMES*, *PPP1R17*), projection neurons (*NFIB*, *SATB2*), inhibitory neurons (*DLX5*, *GAD2*). (F) Triple immunofluorescence for SOX2 (red), TBR2 (green), and MAP2 (magenta), combined with DAPI staining (white), of hCOs day 90. Box indicates a representative neural rosette that is shown at higher magnification; scale bars: 1000 µm and 100 µm. (G) Triple immunofluorescence for HOPX (red), GAD67 (green), and CTIP2 (magenta), combined with DAPI staining (white), of hCOs day 90. Box indicates a representative neural rosette that is shown at higher magnification; scale bars: 1000 µm and 100 µm. (H) Box plot showing prediction score for the identified cell type in scRNA-seq analysis different datasets, split by protocol (guided and unguided protocols), (see Table S1).

To gain insights into lineage dynamics, we performed RNA velocity where we combined cells from 3 developmental timepoints (day 30, day 90, and day 166) (Figures S1D-S1G). This analysis revealed a dynamic transition from neural progenitors to neurons. As observed in organoids from H1-hESCs (Figure 1H), both 30- and 90-day old organoids from H9-hESCs also produced velocity vectors. H9-hESCs derived from day 30 organoids showed evidence of a potential direct neurogenesis, in which neuronal production stems from vRGs via transient progenitor population of IPs (Figure S1E). At day 90, while H9-hESC-derived organoids also show signs of a potential indirect neurogenesis (Figure S1F), as seen in H1-hESCs (Figure 1G). Overall, we observed that the day 90 time point has sufficient representation of human brain-relevant cell types for subsequent disease modelling.

Next, we analyzed the cytoarchitecture of hCOs. At day 90 we observed neural rosettes consisting of SOX2-positive neuronal progenitors tightly packed and radially aligned in a circular ventricular zone (VZ)–like structures (Figures 2F). Some of the SOX2-positive cells were positioned at the outskirts of rosettes (Figures 2F). In addition, both rosettes and cell outskirt cells were also positive for marker HOPX, together suggesting the presence of oRG-like cells (Figures 2F and 2G) ^61^. Further, we detected a second concentric circle, subventricular zone (SVZ)–like, composed of TBR2-positive IPs (marker *EOMES*), as found during fetal neurogenesis. We detected neuronal processes (marker MAP2-positive; Figure 2F) and deep layer neurons (marker CTIP2-positive; Figure 2G). We also observed a portion of GAD67-positive INs, thus confirming the observation of INs by scRNA-seq (Figure 2G).

We compared our guided semi-automated protocol for generating hCOs to previously published protocols by comparing scRNA-seq data from our day 90 hCOs to published data from brain organoids cultured for a similar length of time (Table S1) ^36, 37, 40, 41, 62^. Using Seurat’s prediction score metric ^63^ we were able to quantify each cell’s similarity to each cell type from a reference fetal brain dataset ^47^ (see Methods). The type of each cell was predicted, and each cell was given a score between 0 and 1 for how similar it was to that cell type (1 being the most similar). Prediction score analysis revealed that our hCOs were comparable to other guided brain organoid protocols (Figure 2H, Table S1) ^36, 37, 41^. As expected, the unguided (without addition of patterning factors) brain organoid protocol was least similar to the human fetal brain, especially for excitatory neuronal cell types (Figure 2H) ^40^. This analysis shows the semi-automated hCO protocol resulted in physiologically relevant cell types, similar to published guided brain organoid protocols ^36, 37, 41^, but with an increased throughput.

### Large scale differentiation of human cortical organoids from patient lines carrying 16p11.2 hemideletion and hemiduplication

The semi-automated production of hCOs recapitulates many aspects of human neurodevelopment and day 90 hCOs contain an optimal variety of cell types for modelling neurodevelopmental disorders. To test the ability of this platform for modeling human disease, we differentiated hCOs from 18 different donor patient and control lines carrying a 16p11.2 pathogenic CNV (Figures S2). Out of 18 lines, 4 were controls and the remaining 14 carried CNVs in 16p11.2 (6 hemideletion lines, and 8 hemiduplication lines). The hemideletion lines have similar CNVs (size 534 kbp) encompassing 27 protein coding genes, and the hemiduplication lines contain different CNV sizes (five 740 kbp, two 715 kbp, and one 534 kbp), which encompass up to 30 protein coding genes (Figures S2A). We confirmed the normal karyotype and presence of 16p11.2 CNVs using fluorescent *in situ* hybridization (FISH) and array comparative genomic hybridization (aCGH) for all the lines (Table S2; Figures S2B and S2C).

To profile cell type diversity of diseased hCOs, we performed scRNA-seq on 18 different patient cell lines at day 90 (Figures 3A, S3A-S3D). We split 18 lines into two differentiation batches. To determine the batch variability, two control lines, hESC-H9 and iPSC-8402, were included in both differentiations (Figure S3B). Overall, we profiled 167,958 cells, of which the majority clustered in 9 main cell types (Figures 3A and S3C). First, we noticed that hESC-H9 and iPSC-8402 control lines preserve consistency of cell types across batches of differentiation and technical replicates of the same fixed sample for single cell library construction (Figure 3B and S3D). Together, these data show the semi-automated hCO protocol is highly scalable and reproducible. Next, we used cell prediction scores to assess the quality of organoid cells in both patient and control samples. Prediction scores ranged from 0.6 to 0.8 across different cell types (Figure 3C). Importantly, we did not see a significant difference in cell type prediction scores across genotypes (control, hemideletion, and hemiduplication) (Figure 3C). This indicates a similar cell type fidelity to fetal brain tissue across genotypes.

**Figure 3.**
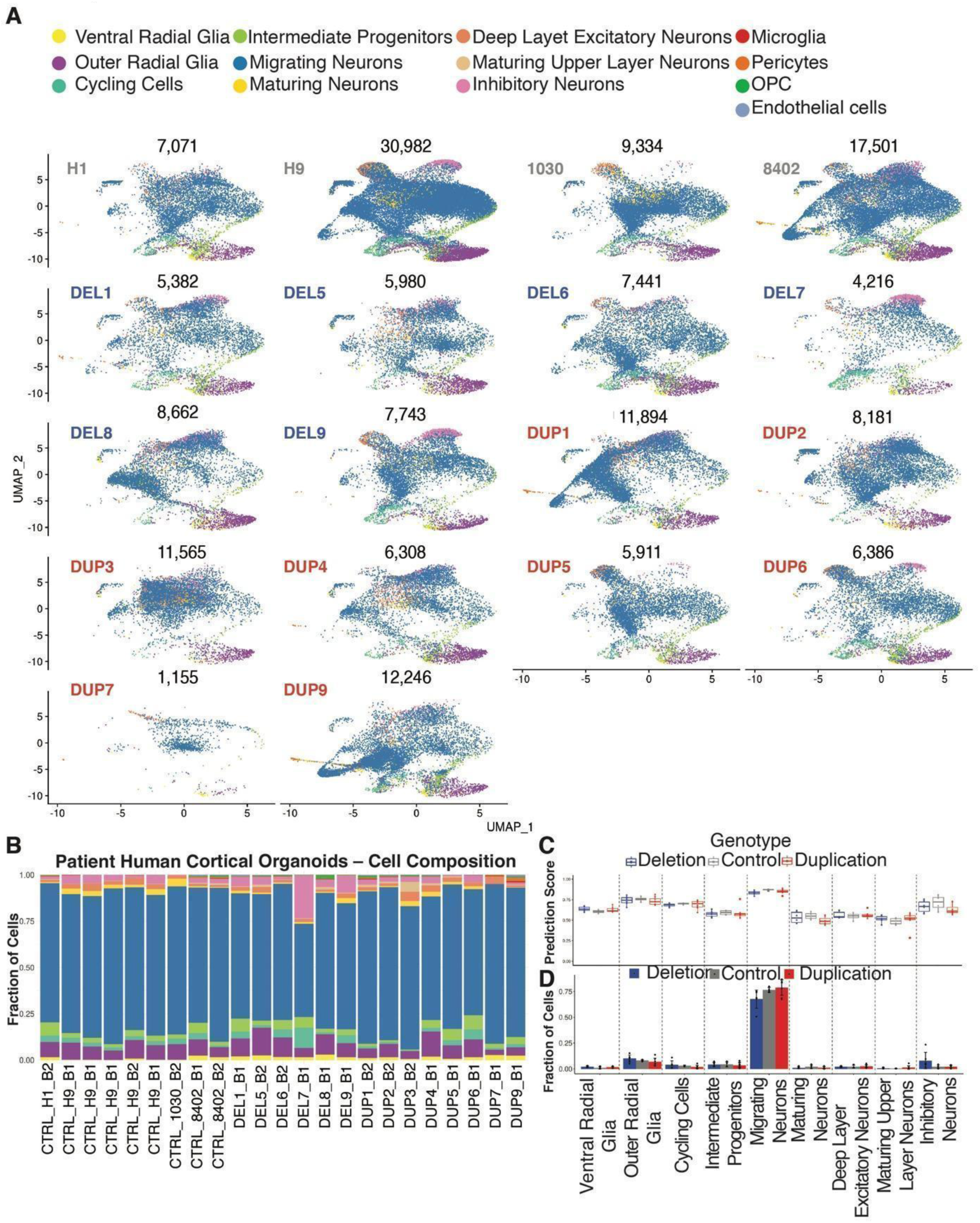
Human Cortical Organoids Derived from Control and 16p11.2 Patient iPSC Lines Show Similar Cell Composition. (A) scRNA-seq data UMAP embedding of individual iPSC lines from day 90-old hCOs after batch correction, control *n* = 4, hemideletions *n* = 6, and hemiduplication *n* = 8; total number of cells 167,958. Note that CTRL_H9 and CTRL_8402 control lines contain higher numbers of cells as they were profiled throughout the multiple 10X single cell runs (see also (see Figure S3D). (B) Stacked bar plots showing relative distribution of cell types measured by scRNA-seq for day 90-old hCOs, each column represents an iPSC line, 10-20 organoids randomly sampled organoids were pulled for each line, note that CTRL_H9 and CTRL_8402 were profiled 5 and 2 times, respectively, in different experiments. (C) Box plot showing prediction score for the identified cell type in scRNA-seq analysis in hCOs, split by genotype, control *n* = 4, hemideletions (DEL) *n* = 6, and hemiduplications (DUP) *n* = 8. (D) Bar plot showing the average fraction of individual cell types per genotype in hCOs, control *n* = 4, hemideletions (DEL) *n* = 6 and hemiduplications (DUP) *n* = 8. Error bars represent standard deviation.

To determine if the presence of the 16p11.2 CNV alters cell composition during development, we quantified the percentage of each cell type across the patient organoids. We did not detect significant differences in cell type composition between the genotypes (Figure 3D). We observed non-significant trends of MigNs decreasing in deletions, and INs increasing in deletions (Figure 3D). Similar fractions of cell types across different donor lines indicate reproducibility of hCOs (Figure 3C and 3D). While we detected some differences between individual donors, when the data was analyzed by genotype, we did not observe a significant alteration in the differentiation propensity for either 16p11.2 hemideletion or hemiduplication lines.

### Cell-type specific alteration of gene expression by 16p11.2 hemideletion

First, to confirm the primary effect of the 16p11.2 CNV, we looked at normalized expression of 16p11.2 gene counts across each genotype. We found significant downregulation of genes in hemideletion donors within the 16p11.2 region (Figure 4A). In contrast, only a few 16p11.2 genes were differentially expressed in hemiduplication donor lines (Figure 4A). Next, we aggregated data from all single cells in each organoid into a pseudobulk transcriptome and performed differential expression across genotypes. From this analysis, we detected ∼50% downregulation of 24/27 genes in 16p11.2 deletion lines (volcano plot genotype averaged, Figure 4B) (see Methods). In addition, we detected multiple differentially expressed (DE) genes outside of 16p11.2 locus (Figure 4C). However, hemiduplications show only a few significantly upregulated, differentially expressed 16p11.2 genes (volcano plot, Figure S4A). This is consistent with the lower penetrance of 16p11.2 hemiduplication relative to hemideletion ^64^. After examining the expression of the 16p11.2 genes, we concluded that the hemideletion samples have a stronger primary deficit and exhibited reduced expression for a higher percentage of the 16p11.2 resident genes in the whole organoid organoid pseudobulk sample compared to hemiduplication lines. At day 90, hemideletions also perturb more genes outside of the 16p11.2 locus indicating a stronger phenotype at this developmental stage. In other studies, the hemideletion consistently produced larger gene expression changes in expression in human lymphoblastoid cells, more severe behavioral phenotypes in mice, and a higher penetrance of phenotypes in humans ^64–67^.

**Figure 4.**
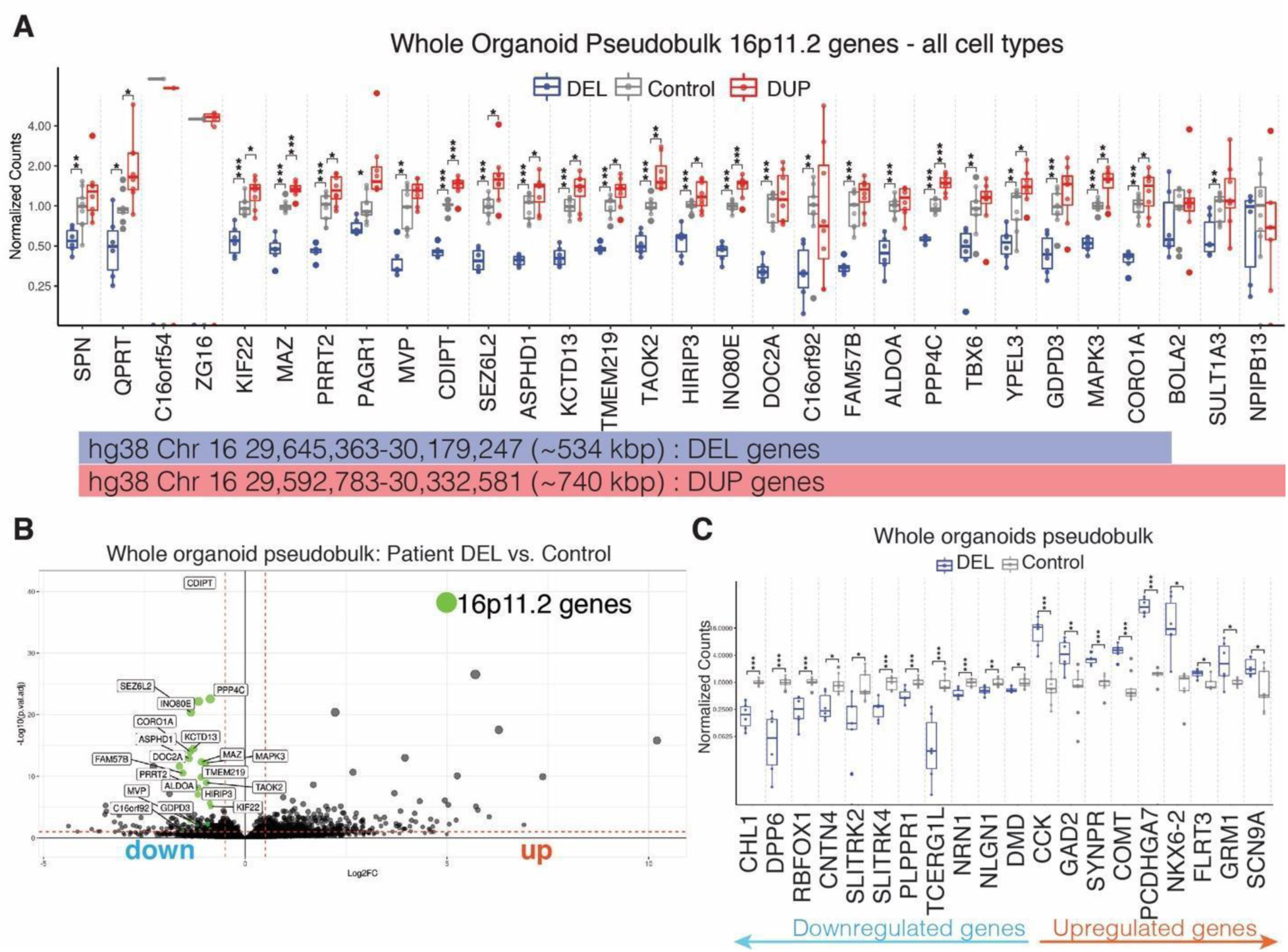
16p11.2 Hemideletion Alters Gene Expression within and outside of Locus in Patients Derived Cortical Organoids. (A) Expression of 16p11.2 genes in hCOs (day 90), pseudobulk analyses in deletion (DEL, blue), control (gray), and duplication (DUP, red); blue and red lane below the graph represent the most common CNV (hemideletion [534 kbp] or hemiduplication [740 kbp]) of the donors analyzed. Each dot presents a cell line. Y- axis represents normalized counts * p < 0.05, ** p < 0.01, *** p < 0.001, (ANOVA test). (B) Volcano plot showing differentially expressed (DE) genes in patient hemideletions vs. control lines, all cell types collectively; 16p11.2 genes are colored green; Cutoff adjusted p-value < 0.1. (C) Selected DEGs that are downregulated and upregulated in patient hemideletion (DEL) vs. control hCOs. * p < 0.05, ** p < 0.01, *** p < 0.001, (ANOVA test).

Next, we examined differential expression across cell types separately for hemideletion and hemiduplication. In hemideletion hCOs, downregulated genes were mostly observed in migrating neurons while upregulated genes were mostly observed in radial glia and migrating neurons (Figures 5, S5A and S5B). In hemideletions we identified 114 DE genes, with 38 downregulated and 76 upregulated across nine cell types (Figure 5, Table S3). Consistent with the relatively low effect on 16p11.2 gene expression, hemiduplication had 70 DE genes, of which 10 were downregulated and 60 were upregulated (Figure 5A). Hemideletion DE genes had overlapping genes across the individual cell types, whereas hemiduplication lines exhibited less overlapping DE genes (Figure 5, Table S3). Notably, in hemideletion hCOs *DPP6, RBFOX1,* and *TCERG1L* were each downregulated in several cell types and are previously associated with ASD and attention-deficit hyperactive disorder (ADHD) (Figure 5) ^68–70^. Furthermore, we found downregulated genes grouped across cell types that play a role in neuronal cell adhesion and were previously associated with ASD including *CHL1*, *CNTN4*, and *NLGN1* ^71–74^ (Figure 5). Interestingly, independent studies showed pathogenic CNVs within *RBFOX1*, *DPP6*, *CNTN4*, *NLGN1,* and *CHL1* in patients suffering from several different neurodevelopmental disorders ^5, 75–77^. Additionally, *COMT* was upregulated in 3 of 9 identified cell types. *COMT is located* in the pathogenic CNV 22q11.2 and was identified as a risk gene for schizophrenia (SCZ) ^78^. Therefore, our analysis of 16p11.2 hemideletion hCOs has identified a potential convergence of 16p11.2 deletion syndrome with other genes previously implicated in neurodevelopmental disorders.

**Figure 5.**
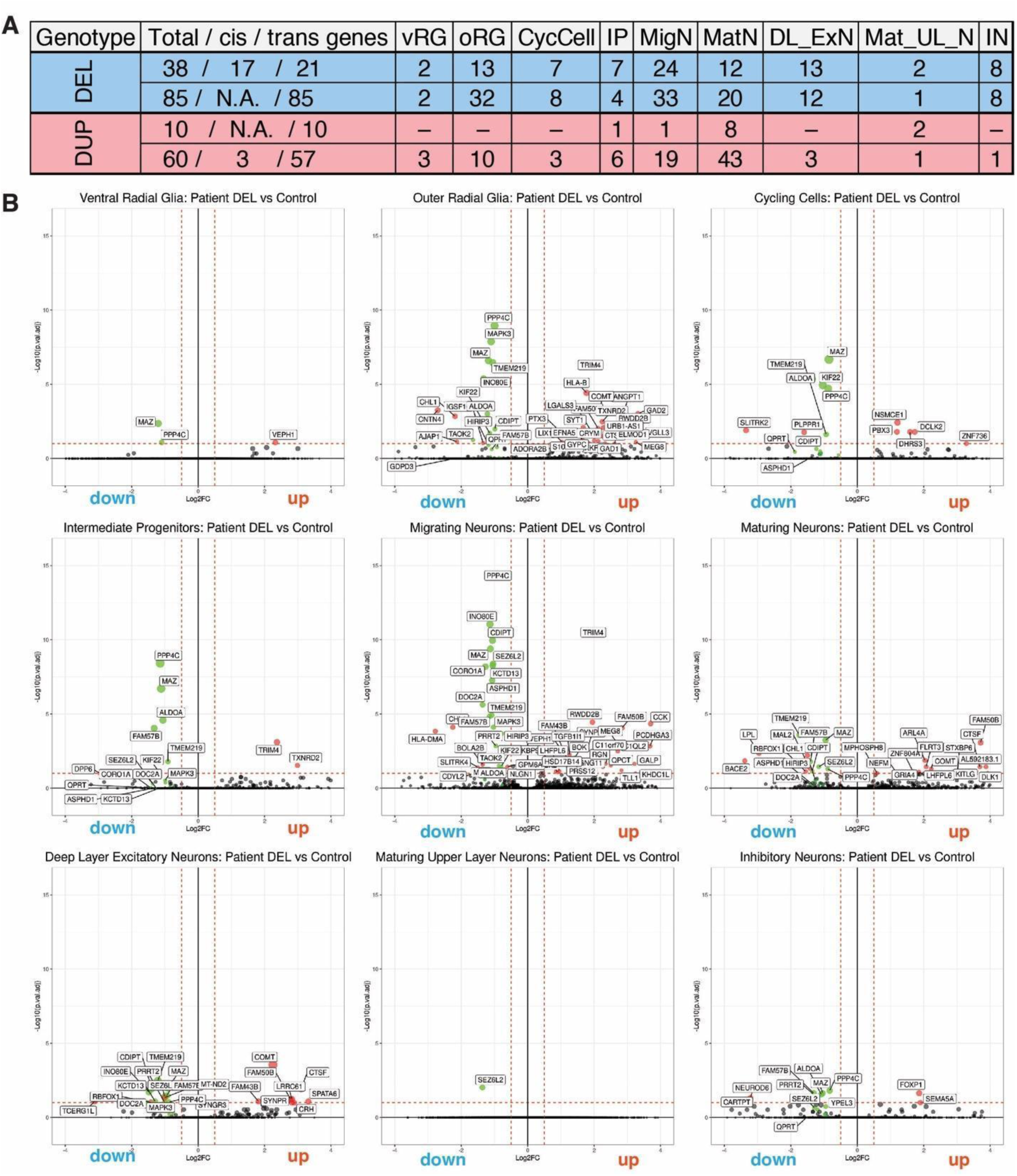
Hemideletion of 16p11.2 Patient hCOs Exhibit Cell-type Specific Alterations of Gene Expression. (A) Table showing number of genes that are downregulated and upregulated in: patient hemideletions vs. control (blue rows), and patient hemiduplication vs. control (red rows), split by the cell type (columns). Cis = genes within 16p locus. Trans = genes outside of the 16p locus. (B) Volcano plots showing DE genes in patient hemideletions (DEL) vs. control lines, split by the cell types; 16p11.2 genes are colored green, selected DEGs are colored red; Cutoff adjusted p-value < 0.1. vRG, ventral radial glia; oRG, outer radial glia; CycCell, Cycling Cells; IP, intermediate progenitors; MigN, Migrating Neurons; MatN, Maturing Neurons; Dl_ExN,Deep Layer Excitatory Neurons; May_Ul_N, Maturing Upper Layer Neurons; IN, inhibitory neurons.

### Pathway and gene–disease network analysis in hemideletion patient lines

Because hemideletion lines exhibit a stronger transcriptional phenotype in our hCO model, we focused our secondary analysis on the hemideletion DE genes. To investigate which processes are enriched across cell type populations, we performed pathway analysis. First, using gene set enrichment analyses (GSEA) we identified 11 different clusters containing gene ontology (GO) and biological processes (BP) IDs (Figures 6A and S6A, Table S4). Next, we designated relevant functional categories for each of these 11 clusters, based on the most common GO IDs (Figure S6A) (see Methods). The most robust enrichment of different functional categories was observed in oRGs, vRGs, IPs, MatNs, and DlExNs (Figure 6A). We detected prominent enrichment of upregulated genes in several neuronal populations in multiple clusters (e.g., response to neuron cell death, biosynthetic processes, synaptic processes, sensory perception, regulation of neurogenesis, and neuronal differentiation). This suggests a change in maturation of excitatory neurons in hemideletion patient lines, similar to observations made using a 2D directed differentiation of hemideletion patients’ iPSCs to neurons ^31^. In addition, highly proliferative oRGs and IPs were enriched in cell cycle, chromosome segregation, and biosynthetic processes (clusters 1,3, 4). Lastly, multiple DE genes in hemideletions were brain-specific genes (Figures S6B and S6C).

**Figure 6.**
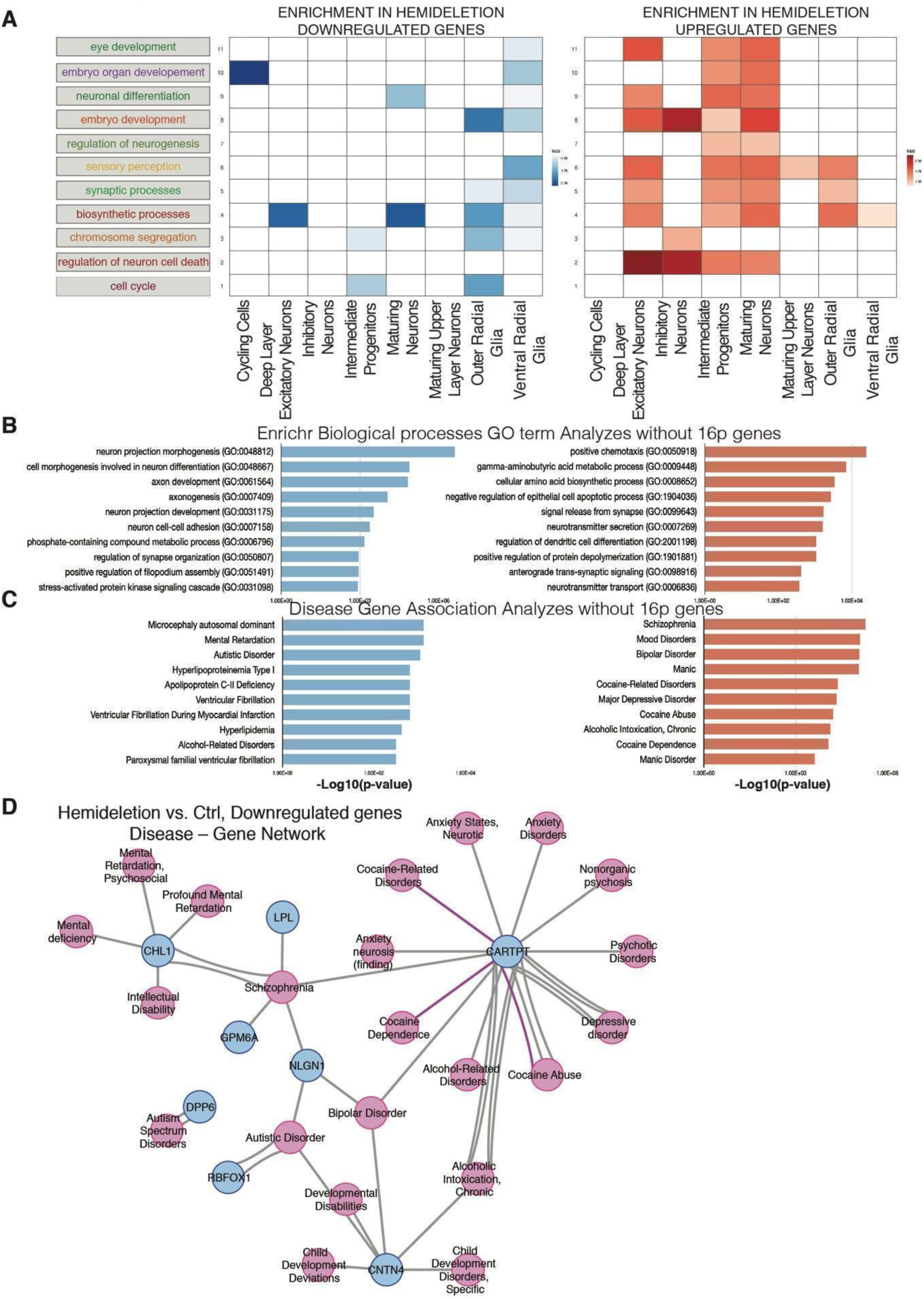
Functional Categories, Gene Ontology, and Disease–gene Enrichment Analysis of Day 90 Cortical Organoids in Control and 16p11.2 Patient Hemideletion Lines. (A) Enrichment of functional categories derived from gene ontology (GO) clustering of genes that are downregulated and upregulated in hemideletions, split by cell types. (NES, normalized enrichment score); (See Table S4) (B) Top 10 GO: biological processes for downregulated (blue) and upregulated (red) genes in hemideletions (cutoff adjusted p-value < 0.1), ordered by their enrichment p-value; x-axis is converted –log10(p-value) (See Table S4). (C) Top 10 gene disease associations, for downregulated (blue) and upregulated (red) genes in hemideletions (cutoff adjusted p-value < 0.1), ordered by their enrichment p-value; x-axis is converted –log10(p-value) (See Table S5). (D) Graphical representation of gene–disease network for downregulated genes in patient hemideletion hCOs (See Table S5).

To specifically elucidate pathways associated with DE genes outside of the 16p11.2 locus, we excluded 16p11.2 genes and performed GO: BP term analysis of downregulated and upregulated genes in hemideletions, using the Enrichr gene analysis platform ^79–81^. Downregulated genes were implicated in neuron projection morphogenesis, neuron cell-cell adhesion, axon development and other terms implicating neurite process assembly and neuron-to-neuron interaction (Figure 6B). This result is consistent with a recent study which identified defects in neuronal migration in 16p11.2 hemideletion hCOs ^82^. Furthermore, upregulated genes were implicated in synaptic-relevant processes such as signal transmission and neurotransmitter biosynthesis and transport (Figure 6B, Table S4). Together, gene enrichment analysis shows signatures of disrupted neuronal maturation and synaptic function in 16p11.2 hemideletion hCOs.

To investigate a direct link between differentially expressed and disease genes, we used the Cytoscape disease–gene network tool ^83, 84^ to perform disease gene enrichment (Figure 6C, Table S5). We found that microcephaly, intellectual disability, and autistic disorder were the top three enrichments for downregulated genes. Moreover, we generated a disease–gene network for downregulated genes, which illustrates that several genes (*RBFOX1*, *CNTN4*, *DPP6*, *CHL1*, and *NLGN1*) converge upon neurodevelopmental and neuropsychiatric disease-gene networks. (Figures 6C and 6D). In addition, analysis of upregulated genes identified enrichments in schizophrenia, mood disorders, and bipolar disorder. Together, disease–gene enrichment analysis suggests that in hemideletion hCOs, differentially expressed genes located outside of the 16p11.2 region are associated with neurodevelopmental disorders.

### 16p11.2 patient hCOs exhibit reduced RBFOX1 protein expression

*RBFOX1* is a critical neuron-specific regulator of alternative splicing in human neurodevelopment ^2^. Similar to 16p11.2 deletions, haploinsufficiency of the region in 16p13.3 containing *RBFOX1* is associated with ASD, epilepsy, and ID ^1^. We found that *RBFOX1* mRNA is downregulated in 16p11.2 hemideletion patient hCOs, specifically within the neuronal cell types (Figures 5B, S7A, and S7B). We reasoned that lower mRNA expression of *RBFOX1* in hemideletion patient lines would result in lower protein expression. To check this, we used SOX2 and RBFOX1 antibodies to perform immunohistochemistry on fixed 20 µm thick hCOs cross-sections. As expected, SOX2 and RBFOX1 showed nuclear localization, in mutually exclusive cell populations (Figure 7A). SOX2-positive nuclei, as seen before (Figure 2F), were labeling neural progenitor rosettes, while RBFOX1-positive nuclei were found in adjacent neuronal populations (Figure 7B). To quantify the intensity of nuclear RBFOX1 and SOX2 fluorescence, we used a high content imager. On average, we analyzed 3 fluorescently labeled cross-sections from 3 independent 90-day old hCOs per cell line (6 control lines [CTRL_H9 batch 1 and 2; CTRL_8402 batch 1 and 2; CTRL_H1; and CTRL_1030] and 6 lines from individual hemideletion patients). Prior to quantification we trained the software to do the following steps: 1) recognize all nuclei using DAPI staining; 2) distinguish live and dead (pyknotic) nuclei based on nuclear size and fluorescent brightness intensity; 3) identify RBFOX1-positive nuclei; and 4) identify SOX2-positive nuclei (Figure 7A). Overall, we measured intensity on 1,096 different fields, containing 1,201,643 live cells, out of which 344,217 nuclei were positive for RBFOX1 (Figure 7A, Table S6). We found that the percentage of RBFOX1-nuclear positive cells was significantly lower in hemideletion lines (Figure 7C). In addition, RBFOX1 nuclear intensity was significantly decreased in hemideletion lines (Figure 7D). In contrast, the percent of SOX2-positive nuclei and SOX2 nuclear intensity were comparable between controls and hemideletions (Figure S7C and S7D). This confirms our earlier observation by scRNA-seq that neural progenitor number is comparable in patient-derived hemideletions and controls, similar to Deshpande et al. 2017 ^31^. We noted variability in nuclear RBFOX1 intensity, which could be due to different patient donors. This is not surprising, given the incomplete penetrance and genetic heterogeneity of neurodevelopmental disorders ^6, 64, 85^.

**Figure 7.**
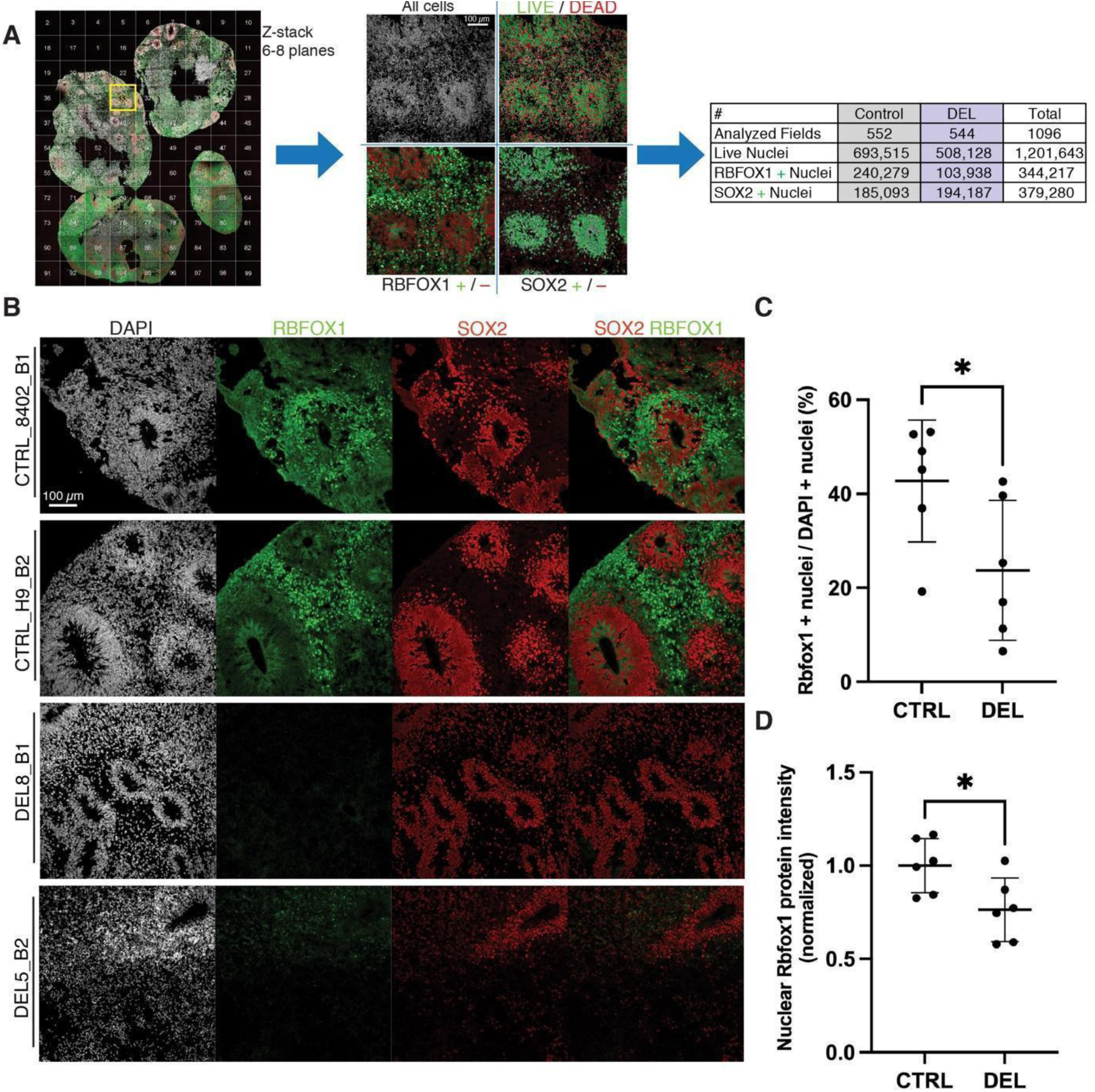
16p11.2 Hemideletion of Human Cortical Organoids Exhibit Reduced Expression of RBFOX1. (A) High content imaging of hCOs cryosections and strategy to detect live cells, RBFOX1-positive cells, and SOX2-positive cells (B) Double immunofluorescence for RBFOX1 (green) and SOX2 (red), combined with DAPI staining (white), of hCOs day 90; selected cell lines CTRL_8402_B1, CTRL_H9_B2, DEL8_B1, DEL5_B2; scale bars: 100 µm. (C) Quantification of the percentage of RBFOX1-positive nuclei over total number of nuclei determined by DAPI on cryosections (20 µm) in control (CTRL, *n* = 6) and hemideletion (DEL, *n* = 6) day 90 old hCOs. Each datapoint presents an independent cell line, with CTRL_H9 and CTRL_8402 differentiated in two different batches. The mean ± SD is shown; * p < 0.05 (*t*-test). (see Table S6). (D) Quantification of nuclear intensity of RBFOX1 fluorescent signal in all live cells (determined by DAPI, see Methods) on cryosections (20 µm) in control (CTRL, *n* = 6) and hemideletions (DEL, *n* = 6) day 90 old hCOs. Average intensity of fluorescent signal value of all controls was used to normalize data. Each datapoint presents an independent cell line, with CTRL_H9 and CTRL_8402 differentiated in two different batches. The mean ± SD is shown; * p < 0.05 (*t*-test), (see Table S6).

After confirming the reduction of RBFOX1 protein in 16p11.2 hemideletion hCOs, we next looked for evidence that a reduction in RBFOX1 could influence the changes in gene expression observed in 16p11.2 hCOs. RBFOX1-dependent splicing events and secondary changes in gene expression have been previously determined by performing RNA-seq after *RBFOX1* knockdown in differentiated primary human neuronal progenitors ^2^. In that study, RBFOX1 regulated 1560 genes which were enriched for regulators of neurodevelopment and autism. By comparing these published RBFOX1 targets to the DE genes we identified in the 16p11.2 hemideletion hCOs by pseudobulk RNA-seq analysis, we found that ∼21% of the DE genes were also regulated by RBFOX1 (24 of 114 genes, adjusted-p < 0.1, whole dataset, pseudobulk; Figure 4B and 4C, Table S7). Some of the 24 genes that overlap between 16p11.2 hCOs and RBFOX1 targets have been previously linked to neurodevelopmental disorders (e.g., *NLGN1*, *CHL1*, *SLITRK2*, *SYNPR, CRH*, *DCLK2*, *TCF7L2*, *GAD1*, *GAD2*, *LGALS3*) ^76, 86–95^. Overall, these data suggest that the reduction in RBFOX1 protein in 16p11.2 hemideletion hCOs is responsible for some of the differential gene expression and indicates the existence of a common molecular signature across independently associated neurodevelopmental risk loci.

## DISCUSSION

This study generated single cell transcriptional profiles from 16p11.2 hemideletion patients and controls in a complex 3D human organoid model, which recapitulates early human brain development *in vitro*. We found that hCOs have transcriptional similarities with the developing fetal neocortex atlas ^47^. hCOs showed a clear maturation trajectory along the neuronal lineage from neural progenitor cells to mature neurons at the oldest profiled age (166 days). In addition, our analysis of lineage relationships suggested an occurrence of crucial events such as direct and indirect neurogenesis ^49–51, 53^. By using automation we were able to standardize and scale-up production to generate hundreds of hCOs to model neurodevelopmental disease using 18 patient and control cell lines. By performing scRNA-seq on over 150,000 cells, we demonstrated similar cellular composition across cell lines and batches of independent differentiations. The robust, large-scale patterned organoid differentiation presented here leveraged automation to produce cells similar to the fetal brain at a sufficient scale to identify a transcriptional signature in 16p11.2 hemideletion. Furthermore, we were able to utilize high throughput quantification of immunofluorescently stained organoid sections to validate the reduction of RBFOX1 at the protein level by examining over a million cells across patients and controls. Overall, this demonstrates that brain organoids at scale will continue to play a critical role in neurodevelopmental disease modelling. In the future, organoids will become increasingly important for drug discovery projects aimed at demonstrating efficacy, specificity, and safety of new treatments in patient cell lines.

By modelling 16p11.2 CNVs with patient-derived hCOs in combination with scRNA-seq, we elucidated changes in the expression of genes previously implicated in neurodevelopmental disorders. Most of the transcriptional changes we found in hemideletion lines were specific to neuronal cell types, and gene enrichment analysis showed neuronal maturation and synaptic function relevant signatures in hemideletion patients. Despite the apparent methodological differences, the overall finding of increased neuronal maturation in hemideletions was similar to a recent study using 3D brain organoids ^82^. We observed minimal changes in hemiduplication lines. There were fewer DE genes and less of an overlap across cell types. More work will be needed to identify relevant phenotypes in duplication lines, and it may require a broader range of developmental time periods and the use of additional phenotypic assays.

Here we identified a shared molecular signature between independent neurodevelopmental risk loci, the 16p11.2 CNV and the *RBFOX1* gene, both of which present a similar spectrum of clinical phenotypes in ASD, ID, SCZ, and epilepsy ^1, 3, 5^. Previous studies indicate three 16p11.2 genes (*KCTD13, HIRIP3, ALDOA*), were found to be RBFOX1-dependent genes, suggesting a reciprocal interaction between the 16p11.2 and RBFOX1 CNV loci. Consistent with this, studies in *Drosophila melanogaster* identified evidence for interactions between 16p11.2 homolog genes and neurodevelopmental genes ^96^. Loviglo and colleagues indicated the 16p11.2 region is a chromatin hub for ASD and detected chromatin contacts between the 16p11.2 region and additional ASD genes such as *PTEN*, *TSC2*, and *CHD1L* ^97–100^. For future studies it will be interesting to determine the molecular determinants of this potential network of 16p11.2 regulated gene expression.

Interestingly, many genes disrupted in 16p11.2 hCOs have also been independently connected to neurodevelopmental disease. Like *RBFOX1*, we also identified differential expression of other notable CNV genes such as *DPP6*, *NLGN1*, *CHL1*, *CNTN4*, and *NLGN1*, which are also associated to neurodevelopmental and neuropsychiatric disorders ^5, 75–77^. Future studies will be required to determine how broadly this spectrum of genetic variants may interconnect phenotypically similar, but genetically distinct clinical cases. Regardless of the outcome, this may inform therapeutic discovery strategies, in terms of whether to focus on common pathways to treat a spectrum of disorders or personalized therapeutics to treat narrow and defined groups of patients. Our work here provides evidence of the former and encourages a direction of rationally synergizing therapeutic discovery between mechanistically overlapping neurodevelopmental disorders.

## FIGURES and LEGENDS

**Figure S1.**
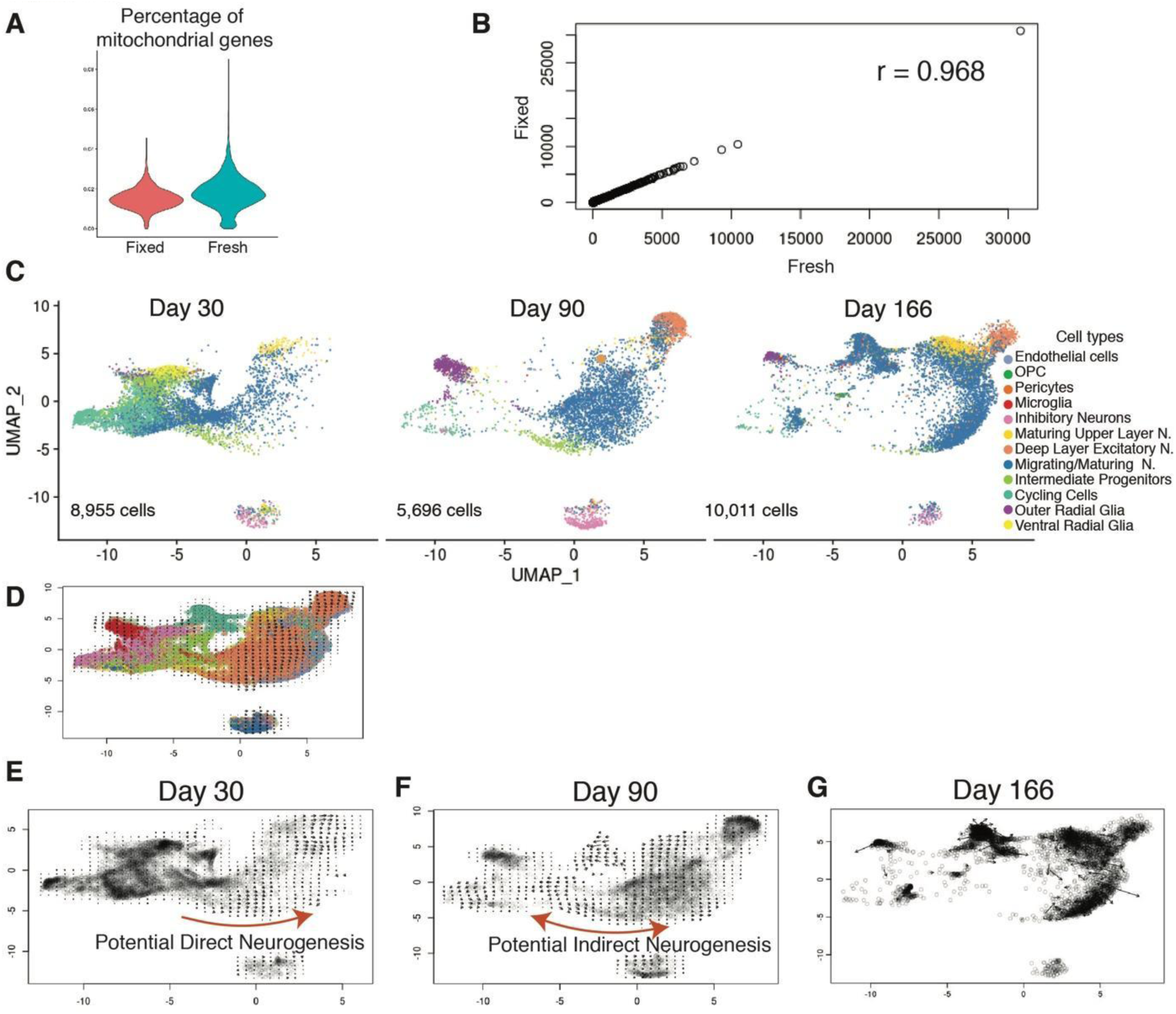
Quality Control of Methanol Fixation Protocol, (A–B) Related to Figure 1; Maturation trajectory of Human Cortical Organoids (C–G) Related to Figure 2. (A) Violin plot showing percentage of mitochondrial genes in fixed and fresh sample (cell line hESC-H1). (B) Paired scatter plot showing correlation of normalized transcriptional count data between fixed and fresh sample (Day 37) (Pearson’s correlation r = 0.968) (cell line hESC-H1). (C) UMAP plot of hCOs scRNA-seq dataset with cell type identities, split by age (Day 30- 8,955 cells, Day 90 - 5,969 cells, and Day 166 - 10,011 cells (cell line hESC-H9). (D–G) RNA velocity of Day 30, Day 90, and Day 166 time points (cell line hESC-H9).

**Figure S2.**
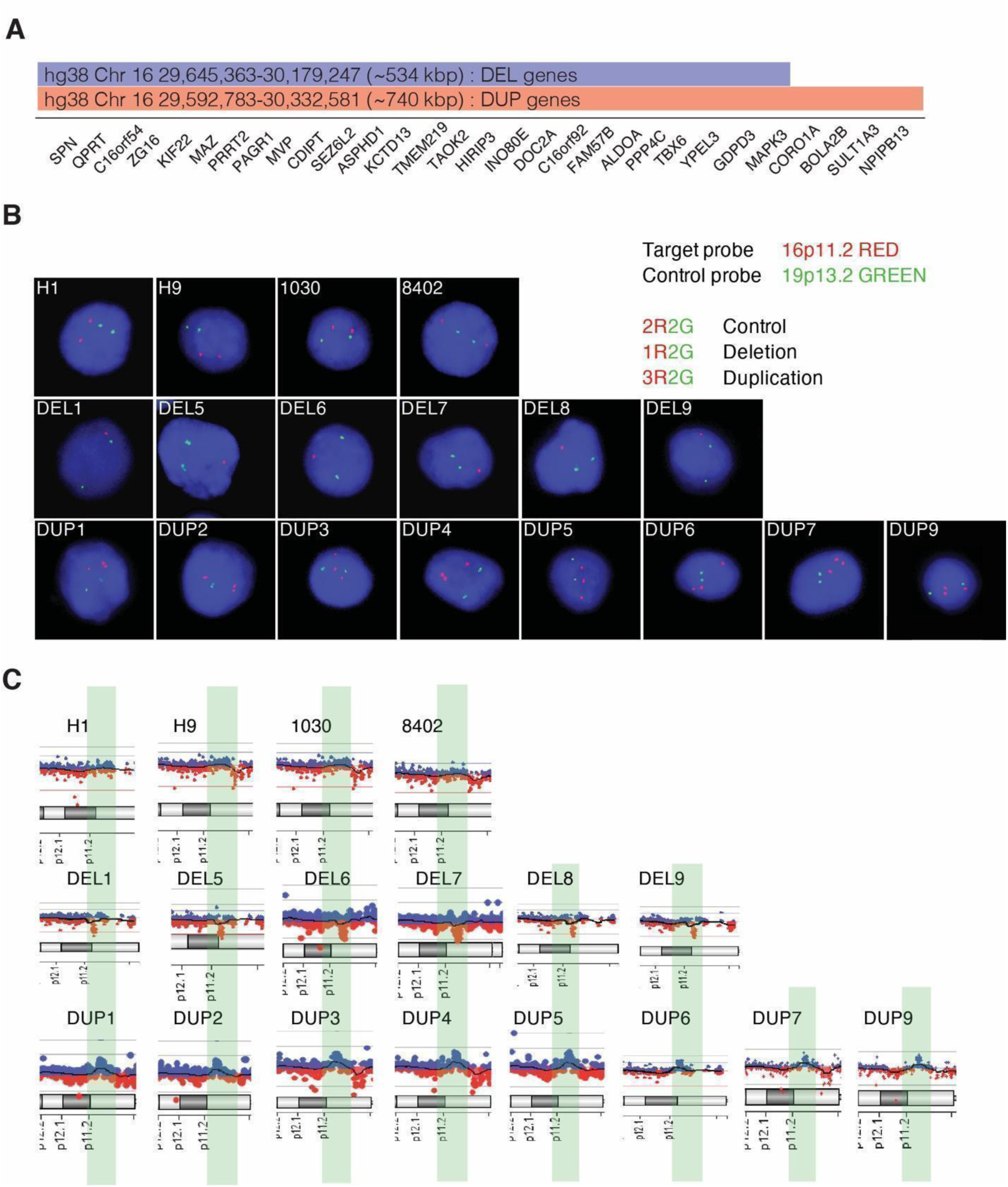
Verification 16p11.2 Hemideletion and Hemiduplication in hPSC Lines Used in this Study, Related to Figure 3. (A) 16p11.2 CNV diagram, hemideletion (blue track), hemiduplication (red track), not to scale. (B) DNA FISH analysis on control *n* = 4, patient 16p11.2 hemideletion *n* = 6, and patient 16p11.2 hemiduplication *n* = 8; nuclei – DAPI (blue), arrowheads pointing at 19p13.2 locus - control probe (green), 16p11.2 locus (red), (see Table S2). (C) Array Comparative Genomic Hybridization (aCGH) on control *n* = 4, patient 16p11.2 hemideletion *n* = 6, and patient 16p11.2 hemiduplication *n* = 8, (see Table S2).

**Figure S3.**
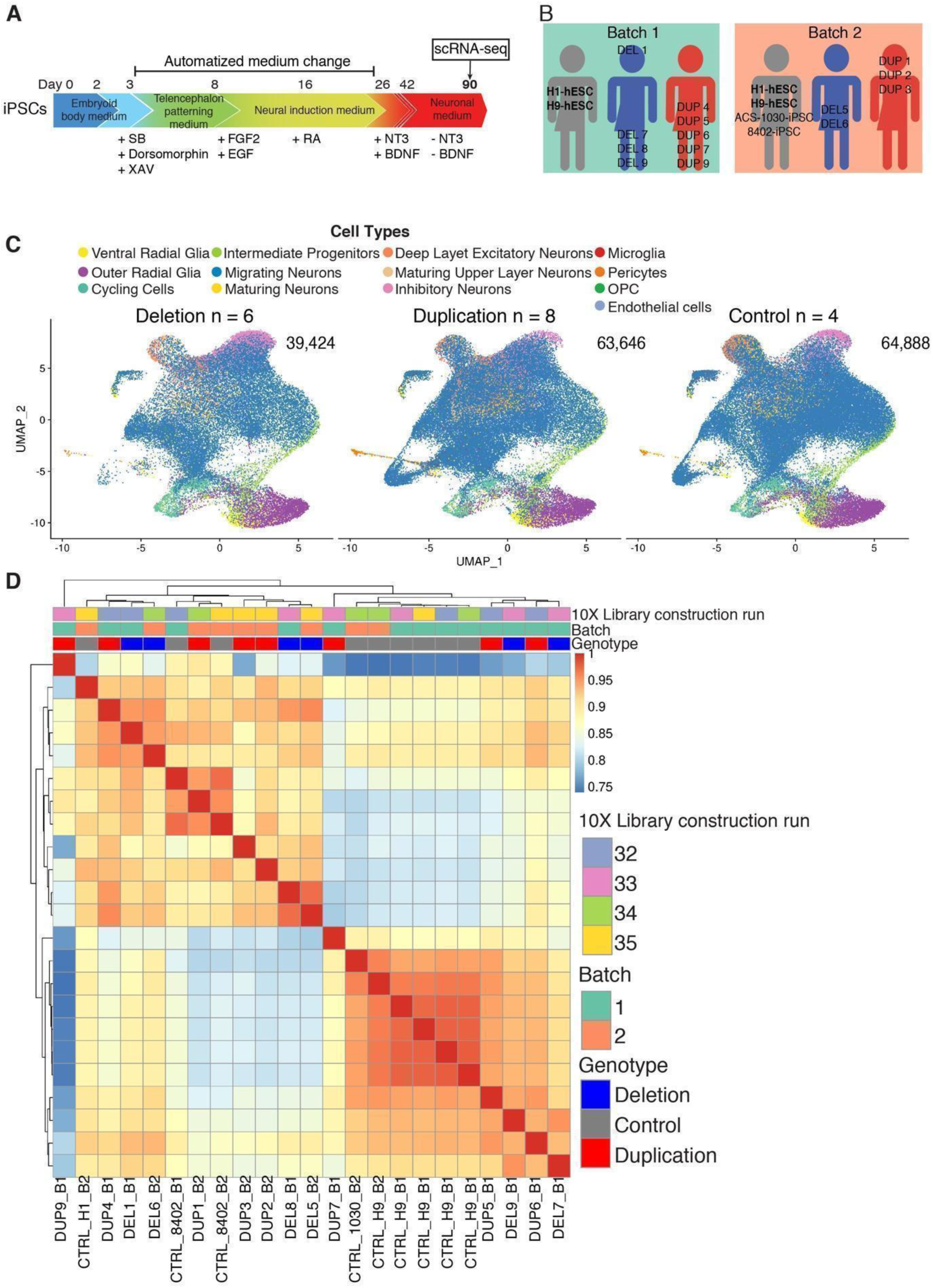
scRNAseq Profiling of Human Cortical Organoids Derived from 16p11.2 CNV Patient Lines, Related to Figure 3. **(A)** Representation of two differentiation batches performed. **(B)** hCOs differentiation diagram. **(C)** UMAP embedding of scRNA-seq data from three-month-old organoids a grouped by sample type to controls (CTRL) *n* = 4, hemideletions (DEL) *n* = 6, and hemiduplications (DUP) *n* = 8; total number of cells 167,958 cells. **(D)** Hierarchical clustering of all samples used in this study.

**Figure S4.**
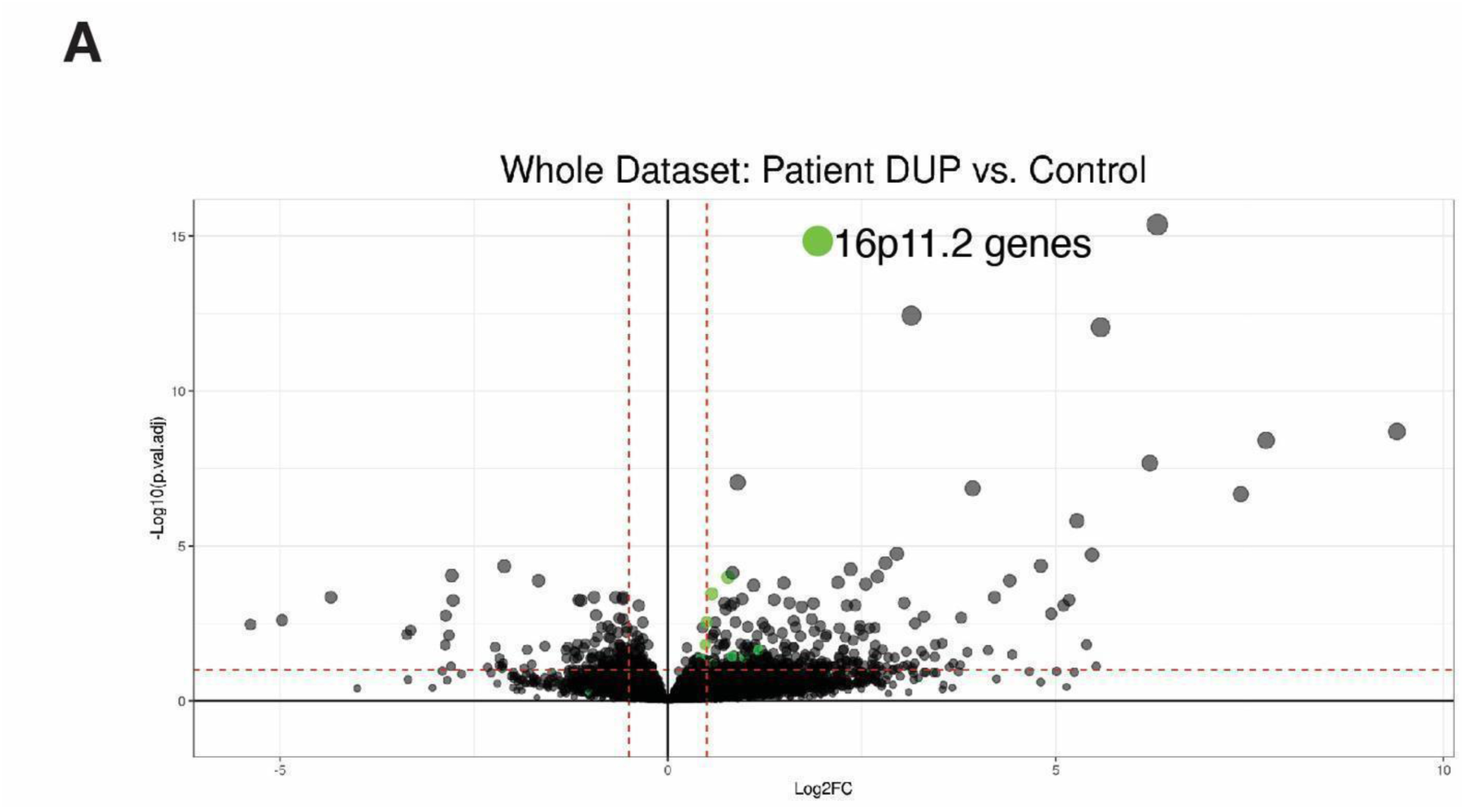
Hemiduplication Patients Derived Human Cortical Organoids do not Exhibit Signature of 16p11.2 genes, Related to Figure 4. (A) Volcano plot showing differentially expressed genes (DEGs) in patient hemiduplication (DUP) vs. control lines, all cell types collectively; 16p11.2 genes are colored green; Cutoff adjusted p-value < 0.1.

**Figure S5.**
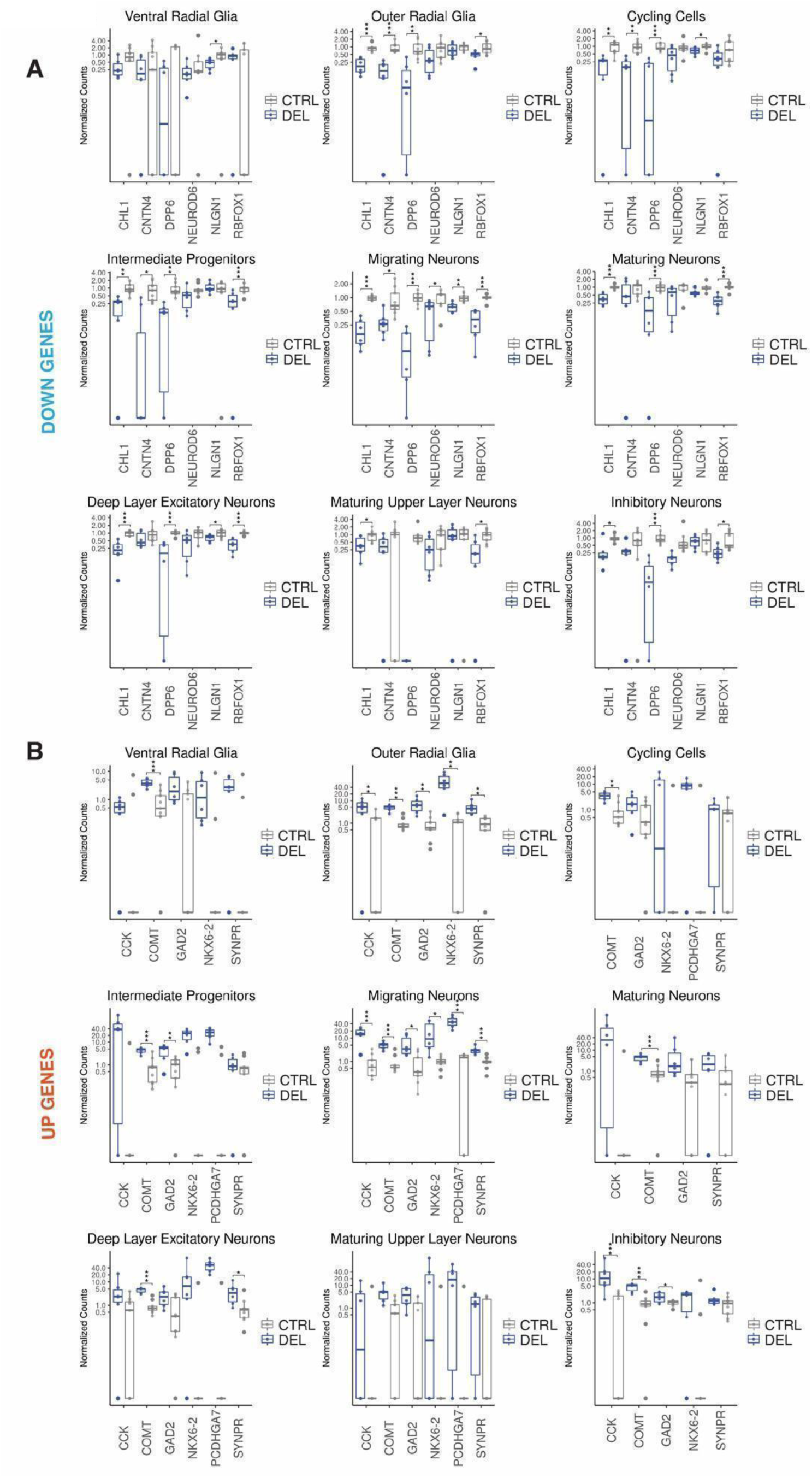
Cell Type Specific Expression of Selected Genes in Hemideletion Human Cortical Organoids, Related to Figure 5. (A) Cell type specific expression of selected genes that are downregulated (blue, vertical) and in patient hemideletion vs. control hCOs. * p < 0.05, ** p < 0.01, *** p < 0.00, (ANOVA test). (B) Cell type specific expression of selected genes that are upregulated (red, vertical) and in patient hemideletion vs. control hCOs. * p < 0.05, ** p < 0.01, *** p < 0.001 (ANOVA test).

**Figure S6.**
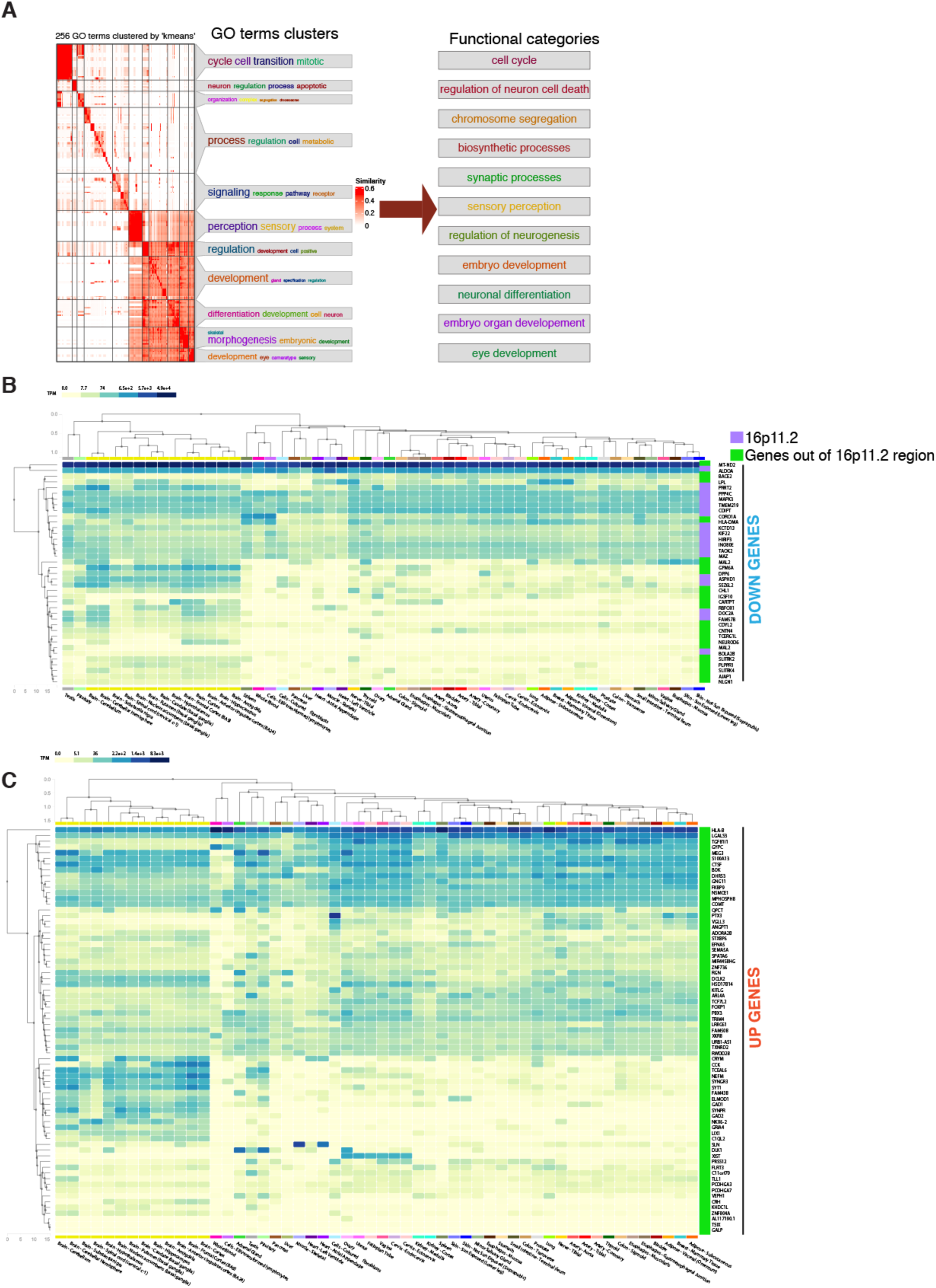
Gene Ontology Analyzes, Related to Figure 6. (A) Gene ontology (GO): biological processes terms clustered by k-means clustering into 11 clusters. GO term clusters were annotated as functional categories into (see Table S4). (B) Heatmap of tissue specific (GTeX) gene query for genes (39) that are downregulated in hemideletion patient derived hCOs. downregulated genes (C) Heatmap of tissue specific (GTeX) gene query for genes (71) that are upregulated in hemideletion patient derived hCOs.

**Figure S7.**
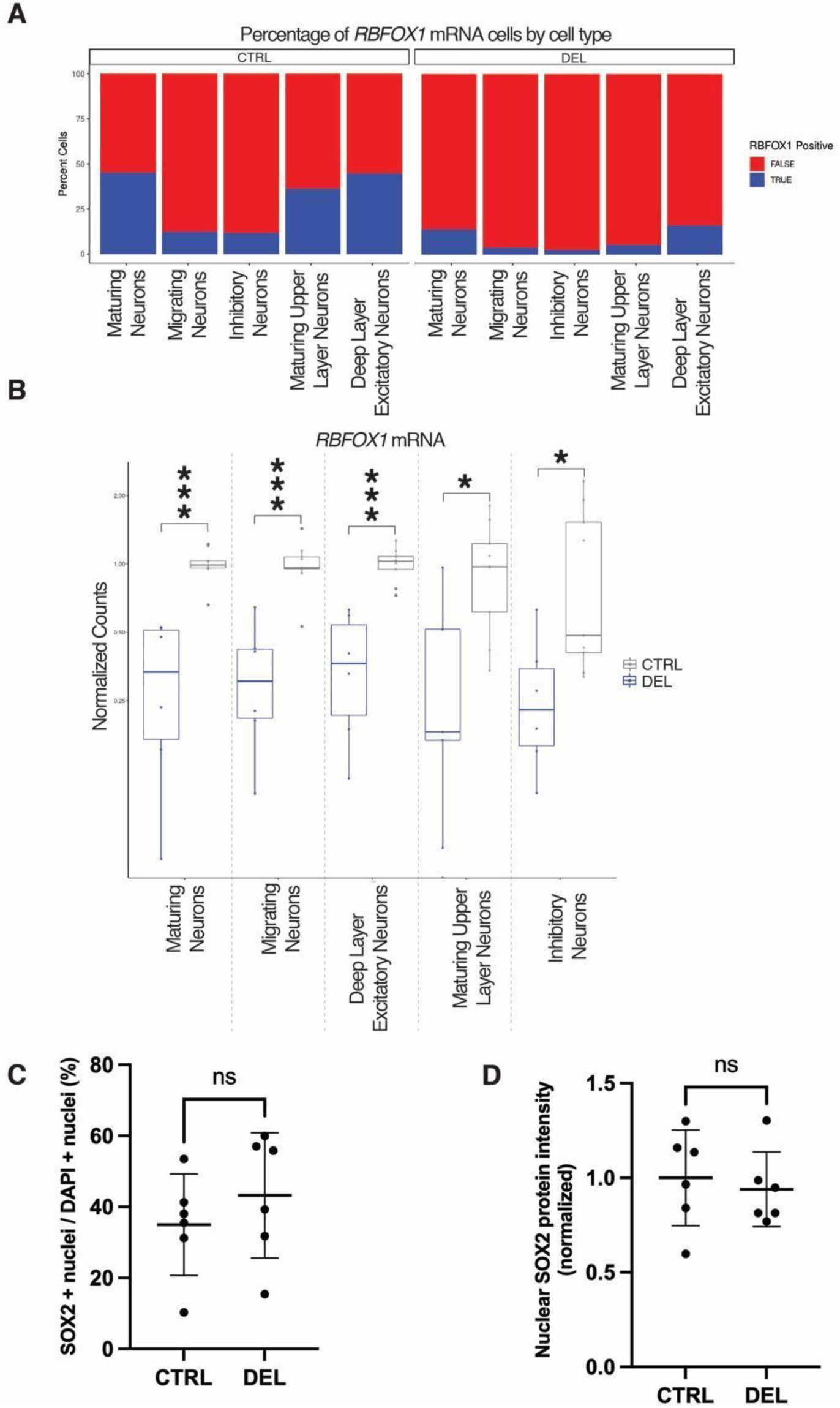
Specificity of *RBFOX1* mRNA Expression, and SOX2 Protein Expression in 90 days old Human Cortical Organoids. Related to Figure 7. (A) Detection of RBFOX1 transcripts in each cell by cell-type in selected neuronal populations in patient DEL vs. control hCOs. (B) *RBFOX1* mRNA expression in selected neuronal populations in patient DEL vs. control hCOs (ANOVA test). (C) Quantification of the percentage of SOX2-positive nuclei over total number of nuclei determined by DAPI on cryosections (20 µm) in control (CTRL, *n* = 6) and hemideletions (DEL, *n* = 6) day 90 old hCOs. Each datapoint presents an independent cell line, with CTRL_H9 and CTRL_8402 differentiated in two different batches. The mean ± SD is shown; * p < 0.05 (*t*-test). (D) Quantification of nuclear intensity of SOX2 fluorescent signal in all live cells (determined by DAPI, see Methods) on cryosections (20 µm) in control (CTRL, *n* = 6) and hemideletions (DEL, *n* = 6) day 90 old hCOs. Average intensity of fluorescent signal value of all controls was used to normalize data. Each datapoint presents an independent cell line, with CTRL_H9 and CTRL_8402 differentiated in two different batches. The mean ± SD is shown; * p < 0.05 (*t*-test).

## List of Supplementary Files

Supplementary Figures S1-S7

## Supplementary tables

Table S1 – List of publications used for benchmarking; Related to Figure 2H.

Table S2 – List of cell lines, metadata, karyotyping, aCGH FISH; Related to Figure S2B and S2C.

Table S3 – List of cell-specific differentially expressed genes (DEGs) in CTRL vs DEL and CTRL vs DUP, filtered to adjusted p-value < 0.1 and unfiltered; Related to Figure 5A-B.

Table S4 – List of GO:BP ids used to designate functional categories; Related to Figure 6A and S6A; List of GO:BP terms formed using downregulated genes and upregulated genes in CTRL vs DEL and CTRL vs DUP, using Enrichr; Related to Figure 6B,

Table S5 – Results of Gene disease network analysis (GDA), for downregulated and upregulated genes in CTRL vs DEL and CTRL vs DUP; list obtained using DisGeNet application in Cytoscape software; Related to Figure 6C and 6D.

Table S6 – Quantification metadata for RBFOX1 and SOX2 percentage and immunofluorescence analysis; Related to Figure 7C, 7D, and Figure S7C, S7D.

Table S7 – List of RBFOX1 dependent genes from (Fogel et al., 2012), list of DE genes from this study (adjusted p-value < 0.1), and overlap of those two lists.

## Methods

### hPSCs lines

A total number of 18 hPSC lines is used in this study. Four control lines, out of which 2 are hESC and 2 hiPSC. Patient-derived iPSCs lines that carry 16p11.2 CNV were split in hemideletions (6) and hemiduplication (8). Each hPSC line represents the individual donor, except two clones lines, DUP3 and DUP 7, which are derived from the same patient. Detailed description of the cell lines is in the supplementary Table S2. Material transfer agreements were arranged with Stanford, NIH and Simons foundation to procure induced pluripotent stem cell lines from consented patients in this study.

### CNVs analyses and karyotyping

To verify CNVs (hemiduplication and hemideletion) in 16P11.2 patient iPSC lines the aCGH tests were performed. Fluorescent *in situ* hybridization (FISH) method was performed to evaluate purity of patient lines. FISH probes were designed to detect signals within chr16:29729098-29909316 in hg38 (UCSC Genome Browser). All lines used in this study were karyotypically normal. aCGH and FISH tests were performed by Cell Line Genetic (Table S2). Karyotyping was performed by Cell Line Genetic or WiCell. See Table S2 for results of the karyotyping, aCGH, and FISH analysis.

### Culture of hPSCs

Feeder free hPSCs were cultured on the 6-well plates (VWR, 10861-554) that were coated with Matrigel (Corning®, 354277, lot no. 9203011) in mTeSR^TM^-Plus (StemCell Technologies, 05825). Cell were passaged, when confluence of 70-80% was reached, using ReLeSR (StemCell Technologies, 05872) to detach and dissociate the colonies, which were then transferred to the mTeSR^TM^-Plus with 0.2 µM thiazovivin added (EZSolution, 1736). All cell lines were karyotypically normal and testing was performed by WiCell and Cell Line Genetics. hPSC lines were tested negative for mycoplasma.

### Cortical organoid differentiation protocol

hPSCs were grown in mTeSR^TM^-Plus. On day 0 cells were inspected under a brightfield microscope to check for differentiation. Cells were used for cortical organoid differentiations when they reached 70-80% of confluence. Prior to lifting the cells, undifferentiated hPCS were primed for 2 hours with 50 µM Y-27632 ROCK inhibitor at 37°C (Selleckchem, S1049), diluted in mTeSR^TM^-Plus. Cells were incubated 6-7 min in Accutase (Gibco^TM^, A1110501) at 37°C. To obtain single cell suspension, 4-5 trituration cycles were performed with P1000 pipette. mTeSR^TM^-Plus with 50 µM Y-27632 was added to neutralize Accutase, cells were transferred to 15 ml tube, spun down 5 min at 200g, resuspended in 5 ml of mTeSR^TM^-Plus with 50 µM Y-27632 and strained through 40 µM strainer (Grainer, 542040). hPSCs’ viability and counts were determined on Vi-CELL XR Cell Viability Analyzer (Beckman Coulter, 731050). To continue the differentiation protocol viability had to be above 80%. Cell were diluted in mTeSR^TM^-Plus with 50 µM Y-27632 to 60,000 cells/ml and 150 µl of cells or ∼9000 cells/well was transferred to ultra-low attachment (ULA) 96-well plates (Greiner, 650979). Lastly, cells were spun 1 min at 150g. On day 2 hPSC embryoid bodies were inspected under the bright field microscope. 120 µl of mTeSR^TM^-Plus with 50 µM Y-27632 medium was replaced with 130 µl of only mTeSR^TM^-Plus medium. From this step on all the feedings of embryoid bodies/cerebral organoids in ULA plates were performed on an automated liquid-handling robot system (Microlab STAR Liquid Handling System, Hamilton Co.). Each cycle of automated feeding performs the following steps: 1) aspirate-dispense 100 µl to lift the dead cell debris, 2) wait 15 s cortical organoid to settle, 3) aspirate 100 µl of old medium, 4) add 100 µl of fresh medium. From day 3 to day 7 cerebral organoids were fed with telencephalon patterning medium (TPM): DMEM/F12, GlutaMAX^TM^ (Gibco^TM^, 10565018), 1x KnockOut^TM^ Serum Replacement (Gibco^TM^, 10828010), 1x Penicillin-Streptomycin (Gibco^TM^, 10378016), 2x GlutaMAX^TM^ (Gibco^TM^, 35050061), 1x MEM Non-Essential Amino Acids Solution (Gibco^TM^, 11140050), 1x 2-Mercaptoethanol (Gibco^TM^, 21985023), 5 µM Dorsomorphin (StemCell Technologies, 72102), 10 µM SB431542 (StemCell Technologies, 72234) and 2 µM XAV939 (StemCell Technologies, 72674). TPM was changed every day from day 3 to day 7. From day 8 to day 25 medium was changed to Neural Induction Medium (NIM): DMEM/F12, GlutaMAX^TM^, and Neurobasal^TM^ (Gibco^TM^, 21103049) in vol/vol 1:1 ratio, 1x B-27^TM^ minus vitamin A (Gibco^TM^, 12587010), 1x N-2 supplement (Gibco^TM^, 17502048), 0.75x GlutaMAX^TM^, 1x Penicillin-Streptomycin-Glutamine, MEM Non-Essential Amino Acids Solution, 1 µg/ml Heparin (Milipore Sigma, H4784), 20 ng/ml EGF (R&D systems, 236-EG-01M) and 20 ng/ml FGF (R&D systems, 233-FB-025) both reconstituted in 0.1% Bovine Serum Albumin. From day 16 to day 25 B-27 without vitamin A is replaced for 1x B-27 with vitamin A (Gibco^TM^, A3582801). On day 26 organoids were transferred to 100 mm petri dishes (Corning^TM^, CLS3262). From day 26 to 42 medium was changed to Neuronal medium that has same base recipe like NIM, but instead EGF and FGF it contains 20 ng/ml BDNF (Peprotech, 450-02) and 20 ng/ml NT3 (Peprotech, 450-03), both reconstituted in 0.1% Bovine Serum Albumin. After day 43 organoids were weaned from BDNF and NT3.

### Cortical organoid dissociation and methanol fixation

Cortical organoid dissociation was performed using the Neural Tissue Dissociation Kit, enzyme (P), (MACS Mitenyi Biotec, 130-092-628). In brief, organoids, 10-30 depending on the size and age, were pooled together and dissociation was performed following manufacturer’s protocol, program 37C_NTDK_1, using gentleMACS^TM^ OctoDissociator with Heaters (MACS Mitenyi Biotec, 130-096-427). Upon dissociation, cells were strained through a 40 µM strainer to achieve single cell suspension. Cells were spun down for 10 min at 300g, supernatant discarded, and cells were resuspended in 1 ml ice cold 1xPBS. Cell number and viability was determined with AO/PI staining on automated cell counter (Nexcelom, Cellometer Auto 2000). To fix the cells, 9 ml of ice-cold 100% methanol was added to the cell suspension in a drop-wise manner to prevent cell clumping. Cells were moved to -20°C for 30 minutes, and subsequently stored to -80°C for long term storage.

### Rehydration of methanol fixed cells

To preserve RNA integrity, it is crucial that all the steps of methanol rehydration were performed on +4°C. Recovery of cells upon methanol rehydration steps was ∼50%. To obtain final suspension of 1×10^6/ml rehydrated cells, 2×10^6 methanol fixed cells were used in each experiment. Cells (2×10^6) were spun down at 4500g for 5 minutes in 4°C precooled centrifuge. The supernatant was removed, and cells were resuspended in 0.5 ml of resuspension buffer: 3xSSC (Milipore Sigma, S6639-1L), 0.04% BSA (Gibco^TM^, 15260037), 1 mM DDT, 0.2 U/µl RNase Inhibitor (Milipore Sigma, 3335402001) in nuclease-free water. Cell were counted with AO/PI staining on automated cell counter (Nexcelom, Cellometer Auto 2000). Finally, dilution of cells was brought to ∼1×10^6 cells/mL, which was the optimal number for downstream single cell sequencing analysis.

### Sectioning of organoids

Organoids were fixed in 4% paraformaldehyde (PFA) in 1 x PBS for 30 minutes at room temperature. Further, organoids were immersed in 30% sucrose solutions in 1x PBS and left on a rocking platform overnight at 4°C. Organoids were embedded in sectioning molds and covered with O.C.T. Compound (Tissue-Tek, 4583) and quickly frozen in ethanol/dry ice bath. Samples were sectioned to 20 µm thickness on Leica CM3050 S cryostat and collected on glass slides. Samples were then stored at -20°C until usage.

### Immunohistochemistry

Samples were thawed at room temperature for 20 min and then washed in 1 x PBS. For staining the nuclear epitopes, an antigen retrieval was performed, except for RBFOX1 nuclear staining (Figure 7), where antigen retrieval was omitted. The antigen retrieval was performed in 1 x Citrate buffer pH 6.0 in ddH2O (Electron Microscopy Sciences, 64142-08). 1 x Citrate buffer pH 6.0 in ddH2O was warmed up to 95°C and applied over samples for 20 min and let it cool at room temperature. Samples were washed 3x with a wash buffer (1 x PBS, 0.1% Tween-20), followed by 30 min permeabilization (0.3% TritonX-100 in 1 x PBS). Samples were quenched with 0.1M Glycine 7.4 pH for 20 min, followed by 30 min incubation in blocking buffer: 5% goat serum (Rockland, D104000050) or 5% normal donkey serum (Milipore Sigma, 566460), 0.1% TritonX-100 in 1x PBS. Primary antibodies were incubated overnight at 4°C, followed by incubation of secondary antibodies for 2 h in the blocking buffer (see Key Resource Table). After two washes in 1 x PBS and one time in ddH20, samples were mounted with VECTASHIELD Vibrance Mounting Medium (Vector Laboratories, H-1700).

### Image acquisition

Images were acquired with an Opera Phenix (PerkinElmer) high-content confocal imager, using a 20x water objective with numerical aperture 1. To set the glass slides adapter was used to position the glass slides into the imager. To locate the specimen (hCOs), a 5x objective was used. Images were acquired in Harmony Phenologic^TM^ (PerkinElmer) software. Each image represents stitched tile scan 41-211 tiles) with 5% overlap, four channels (405, 488, 594, and 647), and 6-8 z-planes. For all images a confocal setting was used.

To stitch tile scans images (Figures 2F and 2G), we exported them out of Harmony Phenologic^TM^ (PerkinElmer) software and imported whole datasets into ArivisVision4D software. ArivisVision4D software allows automatic image stitching. Finally, stitched maximum projections of tiled images were exported as separate .tiff files - 4 different channels (405, 488, 594, and 647). Lastly, final contrast settings and merge of 4 channels was done in ImageJ/Fiji software using functions “B&C” and “merge channels” ^102^. All images represent a maximum projection of 6-8 z-planes of which each z-plane has optical thickness of 1 µm.

### Quantification of immunofluorescence

To quantify nuclear immunofluorescence intensity value on high-content confocal images of hCOs we used “image analysis” mode in Harmony PhenoLOGIC^TM^ software. On average, we analyzed 3 fixed 20 µm thick and fluorescently labeled cross-sections from 3 independent 90 days old hCOs per cell line (6 controls [CTRL_H9 batch 1 and 2; CTRL_8402 batch 1 2, CTRL_H1, CTRL_1030] and 6 cell lines each from individual hemideletion patients). Prior to quantification 6-8 z-planes (each z-plane 2 µm thick) were collapsed into a maximum projection image. We developed custom made pipeline that 1) recognizes all nuclei using DAPI staining; 2) distinguish live and dead (pyknotic) nuclei based on nuclear size and immunofluorescence brightness intensity; 3) determine RBFOX1-positive nuclei; and 4) determine SOX2-positive nuclei. All immunofluorescence signal intensities were kept identical throughout all specimens. Quantifications of selected markers (RBFOX1 and SOX2) in (Figure 7C; Figure S7C) were expressed as a percentage of the number of nuclei for the selected marker over the total number of DAPI-positive live nuclei. Quantification was done on all fields acquired, in total, we measured intensity on 1,096 different fields, containing 1,201,643 live cells out of which 344,217 nuclei were positive for RBFOX1 (∼ 29% out of total live cells) and 379,280 nuclei were positive for SOX2 (∼32% out of total live cells), (Table S6). Intensity of nuclear immunofluorescence signals for selected markers (RBFOX1 and SOX2) were normalized using average intensity of 6 control cell lines (CTRL_H9 batch 1 and 2; CTRL_8402 batch 1 2, CTRL_H1, CTRL_1030) (Figure 7D and Figure S7C).

### Single cell RNA-seq library generation and sequencing

Single cell RNA-seq (scRNA-seq) library construction was performed using the 10X Genomics 3’ Gene Expression kit v3 (cat. #1000075). Rehydrated organoid cells were run through the 10X controller and subsequent library construction was completed in accordance with the manufacturer’s protocol. Library quality was checked using the Bioanalyzer (Agilent Technologies) and sequenced on the NovaSeq 6000 (Illumina). These samples were generated with a target of 4,000-6,000 cells per sample and 50,000 reads per cell.

### Single cell RNA-seq data analysis

Primary analysis of the scRNA-seq data was performed using CellRanger (10X Genomics). BCL files were demultiplexed and converted to fastq format using c*ellranger mkfastq* and fastq files were aligned to HG38 using *cellranger count*. All subsequent analysis was performed in R (v 4.0.2), primarily using Seurat v3 (satijalab.org). Count data from each sample was merged and the aggregated data was normalized, scaled, and technical and biological batch effects were regressed out using SCTransform (Seurat). SCDS was used to help identify and filter out doublets. Cells with a hybrid score greater than 1.4 were removed from the dataset. Dimensionality reduction was performed with principal component analysis (PCA). We determined k-nearest neighbors (KNN) for each cell from this reduction and used the KNN graph to calculate a shared nearest neighbors (SNN) graph. Seurat’s *FindClusters* function was used to identify clusters in the data using a SNN modularity optimization-based clustering algorithm. Cells were labelled using human fetal brain scRNA-seq data as a reference ^47^. The fetal brain raw count data was imported and run through the same basic pipeline as our internal data using *SCTransform* and the original cell type annotations from the authors were used as input for cell type identification in our dataset. Seurat’s *TransferAnchors* function was used to identify fetal brain cell type signatures in the organoid dataset and record each cell’s similarity to those signatures with a score (prediction score) from 0 to 1. Organoid cells were annotated as the cell type with which they had the highest prediction score from the fetal brain data. In Figure 1, internal data and human fetal brain data was integrated using Seurat’s *IntegrateData* function with the human fetal brain data as the reference dataset. RNA velocity analysis was performed using STARsolo (https://github.com/alexdobin/STAR) and velocyto.R (https://github.com/velocyto-team/velocyto.R).

### Pseudobulk differential expression analysis

Pseudobulk analysis was performed by using the sum of all cells from a given sample or all the cells from a defined subset of cells from a given sample as input for differential expression using DESeq2 (https://bioconductor.org/packages/release/bioc/html/DESeq2.html).

### Gene ontology analyses

Gene set enrichment analyses were performed separately on nine cell-types for biological processes (BP) using the clusterProfiler package in R ^103^. The full enrichment tables for the down- and up-regulated genes are shown as heatmaps, in which the color gradients represent the normalized enrichment scores (NESs) (Figure 6A and S6A; Table S4). Next, for an easier interpretation, the enriched gene ontology (GO) terms were clustered based on the similarity matrix of their underlying gene sets using the Bioconductor package simplifyEnrichment ^102^. The resulting 11 clusters were then termed by drawing on the most common words within each cluster.

### Statistical analysis

Statistical analysis was performed either in R (v 4.0.2) using Pearson’s Correlation or ANOVA test (Figures 1D, 4A, 4C, S5A, S5B, S7B) or in GraphPad Prism (v 9.1.2) using *t*-test (Figures 7C, 7D, S7C, S7D).

## Acknowledgements

We are grateful to all the families at the participating Simons Simplex Collection (SSC) sites, as well as the principal investigators at Stanford and NIH who contributed and generated fibroblasts and pluripotent stem cells for this study. We would like to thank Bulent Ataman for thoughtful discussion and feedback on the project.

## Author contributions

M.K., J.J.R., K.A.W., and R.J.I designed experiments and wrote the manuscript. B.H. developed methods to perform automated feeding of organoids. M.K. and Y.S. developed organoid protocols M.K. optimized methanol fixation, and performed IHC analysis. J.J.R performed and analyzed scRNA-seq data. T.T. and J.C. performed gene set enrichment analysis and genomic analysis. J.K. and J.D. cultured organoids. J.S.H. and S.H.C performed copy number analysis and Q.C. of patient iPSC lines. R.D acquired and transferred cell lines, and arranged the MTAs to support this project.

## Data and Code Availability

Raw patient sequencing data will require an MTA from Stanford, Simons Foundation and NIH. Gene count tables and differential expression which lack nucleotide resolution or average expression across patients are available (FILE Sx? GEO?). Code is available via the https://github.com/Novartis (16p11.2scRNA).

## Resources

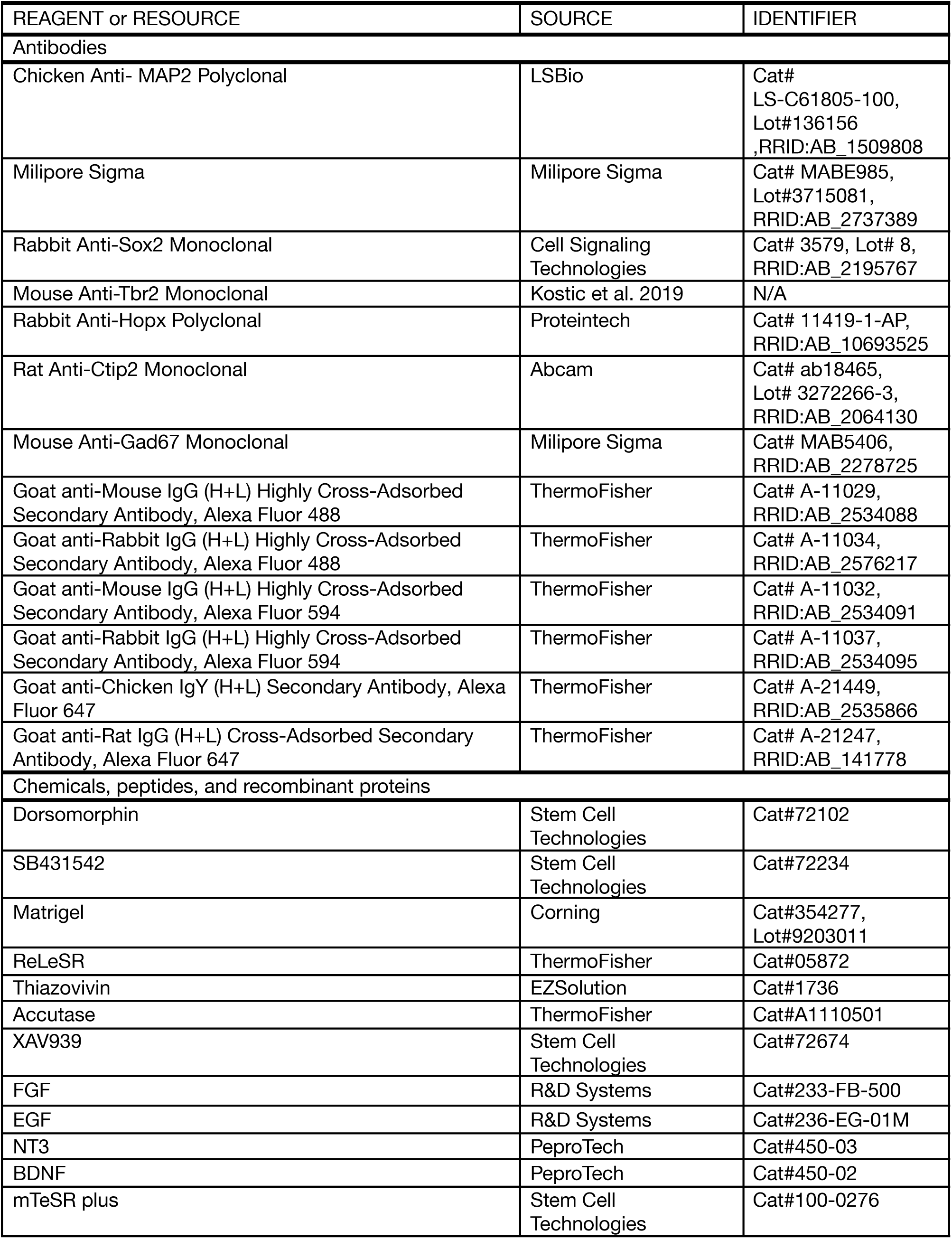
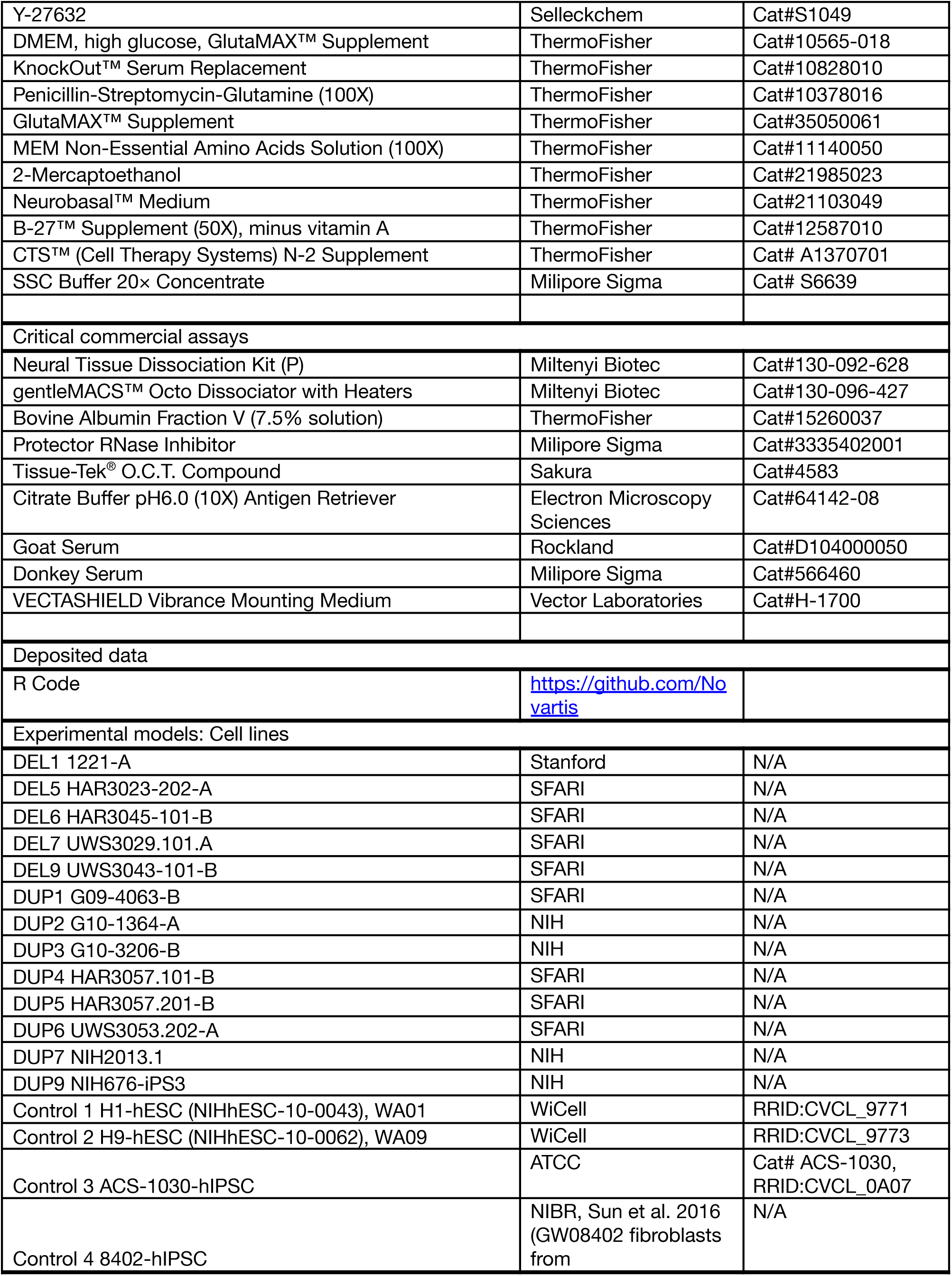
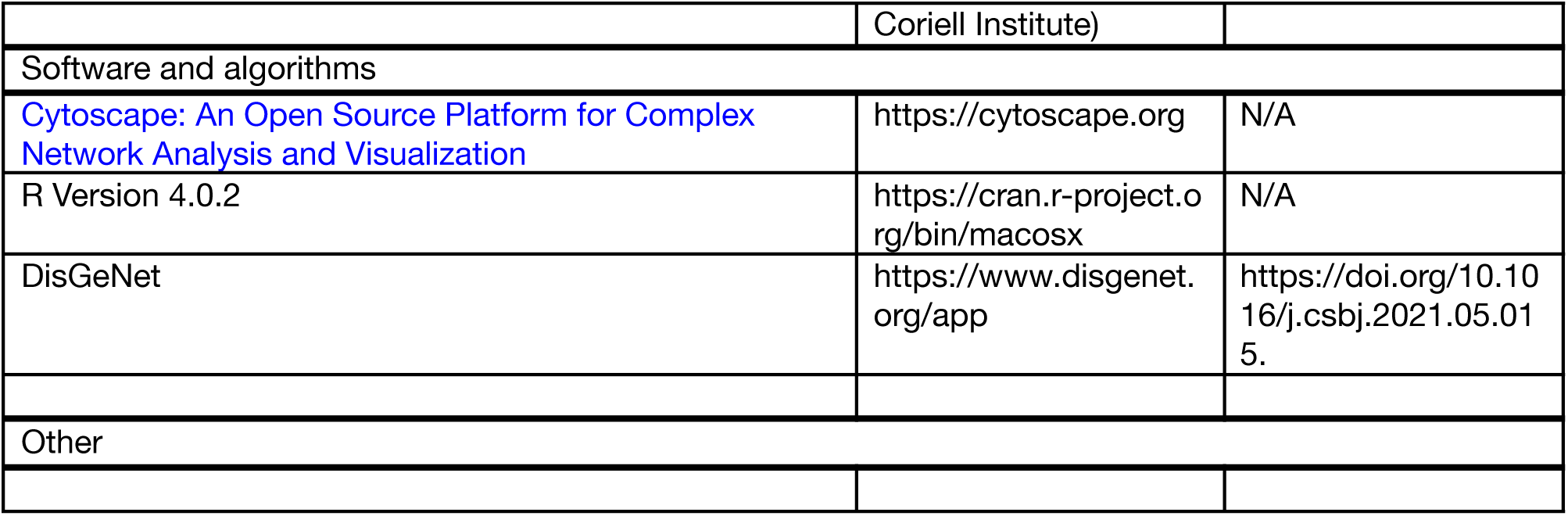

